# Tunneling nanotubes provide a novel route for SARS-CoV-2 spreading between permissive cells and to non-permissive neuronal cells

**DOI:** 10.1101/2021.11.15.468633

**Authors:** Anna Pepe, Stefano Pietropaoli, Matthijn Vos, Giovanna Barba-Spaeth, Chiara Zurzolo

## Abstract

SARS-CoV-2 entry into host cells is mediated by the binding of its spike glycoprotein to the angiotensin-converting enzyme 2 (ACE2) receptor, highly expressed in several organs, but very low in the brain. The mechanism through which SARS-CoV-2 infects neurons is not understood. Tunneling nanotubes (TNTs), actin-based intercellular conduits that connect distant cells, allow the transfer of cargos, including viruses. Here, we explored the neuroinvasive potential of SARS-CoV-2 and whether TNTs are involved in its spreading between cells *in vitro*. We report that neuronal cells, not permissive to SARS-CoV-2 through an exocytosis/endocytosis dependent pathway, can be infected when co-cultured with permissive infected epithelial cells. SARS-CoV-2 induces TNTs formation between permissive cells and exploits this route to spread to uninfected permissive cells in co-culture. Correlative Cryo-electron tomography reveals that SARS-CoV-2 is associated with the plasma membrane of TNTs formed between permissive cells and virus-like vesicular structures are inside TNTs established both between permissive cells and between permissive and non-permissive cells. Our data highlight a potential novel mechanism of SARS-CoV-2 spreading which could serve as route to invade non-permissive cells and potentiate infection in permissive cells.

## Introduction

COVID-19, the disease caused by the severe acute respiratory syndrome coronavirus 2 (SARS-CoV-2), has been developing into a global pandemic since the first reported events in December 2019 (1, 2). Although SARS-CoV-2 primarily targets the respiratory tract and the majority of COVID-19 patients present severe respiratory symptoms (3), other organs such as the intestine, liver, kidneys, heart and brain are also affected. Neurological manifestations of different gravity associated with the COVID-19 have been reported (4–7). Interestingly the neurological symptoms can be acute and resolve with the disease or can represent a major issue in the case of long-COVID (8–10). The ability of SARS-CoV-2 to enter the central nervous system (CNS) is not surprising given that several types of coronavirus (CoV) (e.g., severe acute respiratory syndrome, SARS-CoV; Middle East respiratory syndrome, MERS-CoV) have been reported to invade and persist in the CNS (11, 12). In addition, case reports have shown that the brain tissue of patients that died following COVID-19 were positive for SARS-CoV-2 RNA (13).

However, how SARS-CoV-2 gains access to the CNS and how infection leads to neurological symptoms is still not clear (14–18). SARS-CoV-2 neuroinvasion could be achieved through several routes as previously described (19), and once it reaches the CNS it could bind the angiotensin-converting enzyme 2 (ACE2) receptor exposed on neuronal cells to infect the brain (20). ACE2 receptor is the main actor responsible of the virus entry in the lower respiratory tract (13,21,22). To enter host cells the viral spike (S) proteins of coronaviruses bind the enzymatic domain of the ACE2 receptor, which is exposed on the surface of the cells forming the oral cavity and the oropharynx (23–25). While the expression of the ACE2 receptor has been well documented in many cell type and tissues (23, 24), it is important to underline that in human brain the expression of the ACE2 receptor is low, with the exception of brain areas such as the thalamus and the choroid plexus (26). For this reason, it is not clear how the virus can propagate through the brain, and it is a priority to investigate how SARS-CoV-2 enters into neuronal cells in order to provide new insights in the understanding of the neurological manifestations associated with COVID-19.

An interesting aspect to consider is that the presence of SARS-CoV-2 RNA and proteins has been found in anatomically distinct regions of the brain of COVID-19 patients (13, 27). In this respect, the spreading of SARS-CoV-2 in the CNS is reminiscent of toxic amyloid proteins in neurodegenerative disorders (NDs), that propagate in the brain according to the progression of the pathology (28, 29).

We have previously shown that the spreading of different amyloid aggregates between cells of the CNS and from peripheral lymphoid system cells to neurons occurs mainly through Tunneling Nanotubes (TNTs), a novel mechanism of cell-to-cell communication (30–32). TNTs are thin, membranous conduits rich in actin that form contiguous cytoplasmic bridges between cells over long and short distances (33, 34). Recently we set up a pipeline in correlative cryo-electron microscopy (Cryo-CLEM) to determine the ultrastructure of TNTs in neuronal cell lines, demonstrating their structural identity and differentiating them from other cellular protrusion as filopodia (35). Functionally, TNTs allow direct transport of cargos including virus between distant cells (30,34,36,37). Of specific interest, it has been documented that viruses from different families induce increased formation of TNTs or TNT-like structures, using these membranous structures to efficiently spread the infection to neighboring health cells (38–42). Here, we investigated the neuroinvasive potential of SARS-CoV-2 and whether TNTs are involved in its intercellular spreading. Since TNT-transferred virions would not necessarily be exposed outside the host cell, we hypothesize that TNT-mediated transmission can be used by the viruses to escape neutralizing antibody activity and immune surveillance, as well as to infect less permissive cells lacking the membrane receptor for virus entry, thus allowing for spreading of virus tropism and pathogenicity. Our data show that human neuronal cells, not permissive to SARS-CoV-2 through an exocytosis/endocytosis dependent pathway, can be infected when co-cultured with permissive epithelial cells, previously infected with SARS-CoV-2. We observed, in confocal microscopy, that SARS-CoV-2, induced the formation of TNTs that then could be used by the virus to efficiently spread toward uninfected permissive and non-permissive cells. Furthermore, by setting up correlative Cryo-CLEM and–tomography (35), we demonstrated that SARS-CoV-2 virions are associated to the plasma membrane of TNTs formed between permissive cells. Interestingly, we also observed virus-like vesicular structures and double membrane vesicles (DMVs) inside the TNTs, both between permissive cells and between permissive and non-permissive cells.

Altogether, our results shed new light on the structure of the viral particles undergoing intercellular spreading and provide important information about the molecular mechanism of SARS-CoV-2 infection and transmission. They support the role of TNTs in the viral spreading in both permissive and non-permissive cells, possibly enhancing the efficiency of viral propagation through the body.

## Results

### 1. SARS-CoV-2 can spread among cells through an exocytosis/endocytosis independent pathway

The main route of SARS-CoV-2 entry into the cell is determined by the binding of the spike (S) glycoproteins, exposed on its surface, with the membrane protein ACE2 as an entry receptor (1,24,43,44). Current data indicate that this receptor is only expressed in low amounts in the brain, so the question is whether and how the virus is able to infect neuronal cells (45). To this aim we tested different cell types to verify whether they were permissive to viral infection by the receptor-mediated pathway.

Different cells of mammalian and human origin were plated on a 96 multi-well plate and infected with a MOI (multiplicity of infection) ranging from 10^-1^ to 10^-5^. The cell-lines used in this assay included human colon epithelial cell-lines (Caco-2,), monkey kidney epithelial cell-line (Vero E6), and the human (SH-SY5Y) and murine (CAD) neuronal cells lines. After 3 days we looked for productive infection by staining the infected monolayers with a virus-specific antibody and by titrating the virus released in the supernatant. Using these two parameters we found that only the epithelial Vero E6 and Caco-2 cells were susceptible to infection with SARS-CoV-2, as previously shown (46), while both neuronal cell lines (mouse and human) did not show any sign of infection or viral production (fig. S1A, B). Consistently, the immunofluorescent signal of the viral Nucleoprotein (N) protein was evident in the monolayers of Vero E6 and Caco-2 cells after 3 days of infection with different MOI. On the contrary, no fluorescent signal was detected in CAD, and SH-SY5Y cells (fig. S1A). In addition, an aliquot of the supernatant from the MOI 0.1 infection was collected at day 2 and 3 to quantify the kinetic of virus production by titration using a semisolid plaque assay on Vero E6 cells. As shown in fig. S1 B, no viral production was detected using the supernatant derived from CAD and SH-SY5Y infection, while clear signs of cytopathic effect (CPE) were detected in Vero E6 cells infected with Vero E6 and Caco-2 supernatants.

These data show that neuronal cells cannot be infected directly from the supernatant through a receptor-mediated mechanism.

The main mediator of cellular entry of SARS-CoV-2 is the ACE2 receptor (1,24,47); whilst it is highly expressed in vascular endothelial cells of the lungs (47), it was detected at extremely low levels in the neuronal cells (45). Consistently, we were unable to detect a signal for ACE2 in SH-SY5Y cells (fig. S2), confirming previous observation reporting extremely low levels of expression in neuronal cells (45). Nonetheless, emerging case reports showed that patients infected with SARS-CoV-2 have common neurological manifestation (17,48–53) suggesting that the virus could invade and infect the CNS (7,53,54). Since the ACE2 receptor is not widely expressed in neuronal cells one likely possibility is that the virus can exploit intercellular communication pathways to enter neuronal cells directly from permissive cells, which would allow bypassing the main receptor mediated pathway. To investigate this, we set up co-culture experiments between permissive Vero E6 cells, routinely used as infection model of SARS-CoV-2, and non-permissive SH-SY5Y human neuronal cells. Vero E6 cells (donor cells) infected with SARS-CoV-2 at MOI 0.05 for 48 hours were co-cultured with SH-SY5Y cells (acceptor cells) previously transfected with a plasmid encoding mCherry, to be easily distinguished from donor cells (fig. S3A). After 24h and 48h of co-culture, cells were fixed and immunostained with anti-N antibody recognising SARS-CoV2 nucleoproteins, and labelled with cell mask blue to stain the whole cells (Fig. 1A-C). By using confocal microscopy, and the ICY software (*icy.bioimageanalysis.org*), we calculated the percentage of SH-SY5Y acceptor cells positive for the anti-N antibody immunostaining. After 24h of co-culture, 36,4% of acceptor cells contained in their cytoplasm spots recognised by the anti-N antibody (Fig. 1D), and this percentage increased to 62,5% after 48h (Fig. 1A-D). To further investigate the nature of the viral particles, present in acceptor neuronal cells, additional co-cultures between Vero E6 infected with SARS-CoV2 MOI of 0.05 and SH-SY5Y neuronal cells were immunostained (after 24h and 48h of co-culture) using an anti-Spike (anti-S) antibody, both alone (Fig. 1E-F) and in combination with the anti-N antibody (Fig. 1H-J). Similar to the results obtained with the N-antibody, we found that after 24h of co-culture 21,8 % of acceptor cells contained in their cytoplasm spots positive for the anti-S antibody, and after 48h of co-culture this value increased to 42,4 % (Fig. 1G). We then evaluated the co-localization of the two viral proteins N and S in the acceptor cells (Fig. 1H-J) and we found that at 48h the PCC (Pearson’s coefficient) was in average 0,716, indicating that the two proteins partially colocalize. While separate signals for anti-N and anti-S antibodies could be suggestive of virus uncoating during the first step of the infection (56), colocalization of N and S proteins could correspond to mature virions inside endocytic vesicles that are entering the cell and/or to newly synthetized virions assembled in the neuronal acceptor cells. Because at 24 hours we found negligible colocalization between the two proteins, we could assume that the particles labelled with both the N and S antibodies in the neuronal cells at 48 hours correspond to virus newly assembled in the SH-SY5Y cells after transfer. Indeed at 24 hours of co-culture Vero E6 cell are already full of newly synthetized virions (fig. S3 B, C), therefore if there was direct transfer of fully assembled virions we would have expected to find them in acceptor cells already at 24h of co-culture, which was not the case.

**Fig. 1.**
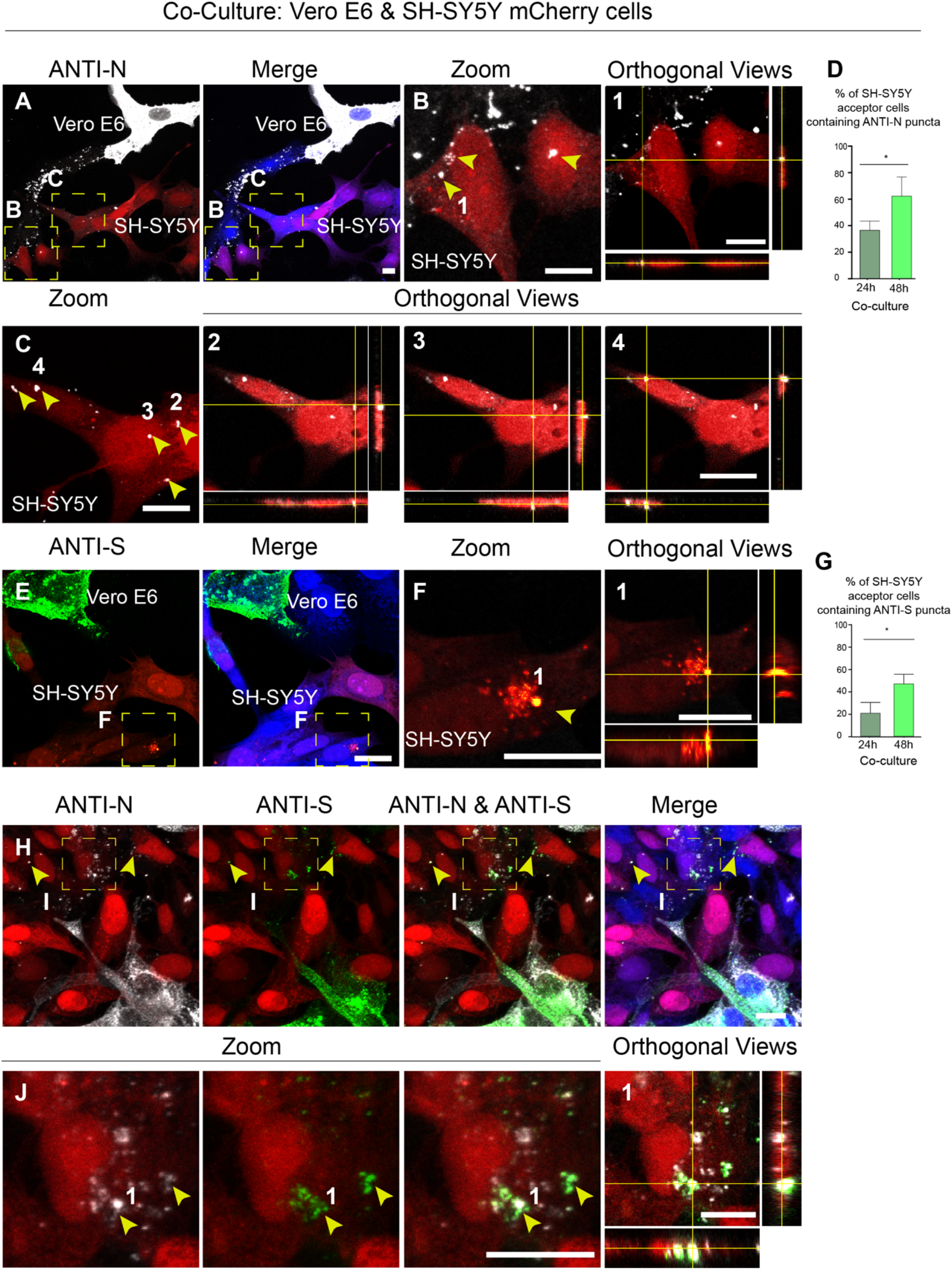
SARS-CoV-2 can reach SH-SY5Y neuronal cells from Vero E6 permissive cells. **(A-D)** Infected Vero E6 cells (donor cells) were co-cultured at 1:1 ratio with SH-SY5Y neuronal cells previously stably transfected with a vector that expresses mCherry (acceptor cells). Co-culture were fixed by using 4% PFA at 24h and 48h. (**A**) Confocal micrographs showing 48h co-culture between SARS-CoV-2 Vero E6 infected cells and SH-SY5Y mCherry cells. Anti-N antibody was used to detect SARS-CoV-2 nucleoproteins; cellular membranes were labelled with cell mask blue. (**B-C**) Enlargement of the yellow dashed squares in (A), the yellow arrowheads indicate the anti-N puncta detected in the cytoplasm of acceptor cells. Numbers (**1, 2, 3, 4**) are the orthogonal views of (B-C) showing the anti-N puncta inside the cytoplasm of acceptor cells. (**D**) Graph showing the mean percentage of N puncta transferred to acceptor cells after 24h and 48h of co-culture: 36.47% ± 3.96 and 62.56% ± 8.28 respectively, (*p=0.0468 co-culture 48h versus co-culture 24h; N=3). (**E**) Confocal micrographs showing 48h co-culture between SARS-CoV-2 cells Vero-E6 infected and SH-SY5Y mCherry cells. Anti-Spike (anti-S) antibody was used to detect SARS-CoV-2 particles; cellular membranes were labelled with cell mask blue. (**F**) Enlargement of the yellow dashed square in (E), the yellow arrowhead indicates the anti-S puncta in the acceptor cells; Number (**1**) is the orthogonal views of (F) showing the anti-S puncta inside acceptor cells. (**G**) Graph showing the mean percentage of S puncta transferred to acceptor cells after 24h and 48h of 24h and 48h: 21.84% ± 5.09 and 42.44% ± 4.38 respectively (*p=0.0374 co-culture 48h versus co-culture 24h; N=3) (**H-J**) Double immunostaining of co-culture using anti-S and anti-N antibodies. (**J**) Enlargement of the yellow dashed square in (H) showing colocalization between anti-N and anti-S puncta in SH-SY5Y mCherry acceptor cells. The Pearson’s coefficient (PCC) between anti-S and anti-N was in average 0,716 (20 cells analysed). Scale bars: A-J 10 µm.

To directly investigate whether SARS-CoV-2 transferred from the donor cells was still functional, and able to replicate in neuronal cells, we performed an immunostaining using the anti-dsRNA (double-stranded RNA) antibody J2 that is the gold standard for the detection of dsRNA in infected cells (57). After 48h of co-culture (Vero E6 infected cells as donors and SH-SY5Y cells as acceptors), cells were fixed and immunostained to detect dsRNA and SARS-CoV-2 particles using the anti-S antibody. We found J2 positive signal both in donor Vero E6 infected cells and acceptor SH-SY5Y cells (Fig. 2A-C). Interestingly, J2 signal in SH-SY5Y cells (Fig. 2B, C) corresponds to bright foci of different sizes and intensities, that could represent the replication organelles (RO) or double-membrane vesicle (DMVs) where viral RNA synthesis occurs (58, 59). Furthermore, the localization of J2 spots in the perinuclear/ER region of acceptor cells could suggest an active viral replication occurring in the neuronal cells (Fig. 2B, C) (60–64). Accordingly, J2 signal was specific for infected cells in co-culture as it was not detected in non-infected Vero E6 co-cultured with SH-SY5Y mCherry (Fig. 2D), as well as, in the negative control in which the co-culture was incubated only with the secondary antibody (Fig. 4E). These data indicate that SARS-CoV-2 transferred by a cell-to-cell contact-dependent mechanism could be actively replicated in neuronal acceptor cells.

**Fig. 2.**
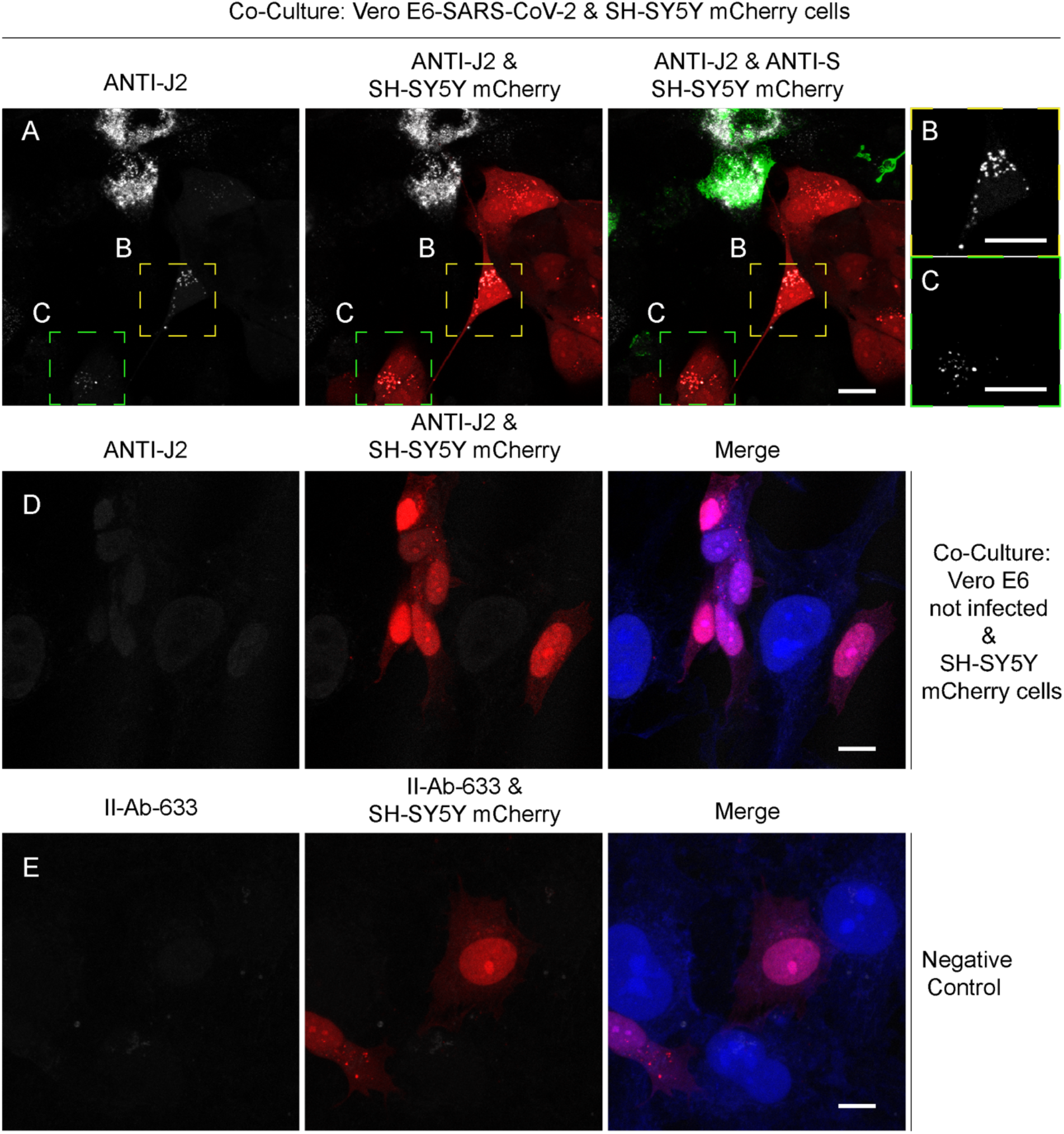
Anti-dsRNA (double-stranded RNA) antibody J2 is detected in SH-SY5Y cells co-cultured with SARS-CoV-2 Vero E6 infected cells. **(A)** SARS-CoV-2 infected Vero E6 cells (donor cells) were co-cultured for 48h with SH-SY5Y mCherry acceptor cells. Confocal micrographs showing the staining with anti-J2 antibody used to detect dsRNA and an anti-S antibody is used to detect SARS-CoV-2 particles. (**B**, **C**) Enlargements of the yellow and green dashed squares in A showing J2 signal detected in acceptor cells. (**D**) Confocal micrographs showing not infected Vero E6 cells co-cultured with SH-SY5Y mCherry cells. The co-culture was immunostained with anti-J2 antibody and cell mask blue. (**E**) Negative control. The co-culture was immunostained with the secondary antibody conjugated with 633 fluorochrome and cell mask blue. Scale bars, 10 µm.

To support this hypothesis, as complementary approach we performed an immunostaining against the non-structural protein 3 (nsp3), an essential component of the viral replication/transcription complex. After 48h of co-culture (Vero E6 infected cells as donors and SH-SY5Y cells as acceptors), cells were fixed and immunostained to detect nsp3 and SARS-CoV-2 particles (by anti-N). We found cytoplasmic accumulations containing the viral N protein and the non-structural protein nsp3 both in donor Vero E6 and acceptor SH-SY5Y cells (Fig. 3A-D). This data confirms that SARS-CoV-2 can replicate in the acceptor neuronal cells. The above data show that neuronal cells can be infected when in co-culture with permissive cells, but do not address the mechanism. Although our data show that SH-SY5Y cells cannot be infected by the supernatant (fig S1), in order to investigate the possible contribution of the endocytic entry pathway in co-culture conditions, as control for the co-culture experiments, we performed “secretion tests” were the supernatants from Vero E6 infected cells were used to infect SH-SY5Y cells for 24h and 48h respectively (fig. S4A). We did not detect any signal for anti-N and anti-S antibodies in the acceptor cells that received the supernatants from the infected Vero E6 cells after 24h (fig. S4B, C); and a negligible signal was found at 48h (fig. S4C). On the other end, the supernatants from Vero E6 infected cells was able to infect control Vero E6 cells where we could detect both anti-N and anti-S signal (fig. S4D). The infectious titer of the supernatant used in the secretion experiment in Vero E6 acceptor cells was calculated using a focus forming assay (fig. S4E). Additionally, the 48h supernatants from donor infected cells, co-culture and secretion experiments were used to assess viral production by focus forming assay titration protocol (fig. S4F).

**Fig. 3.**
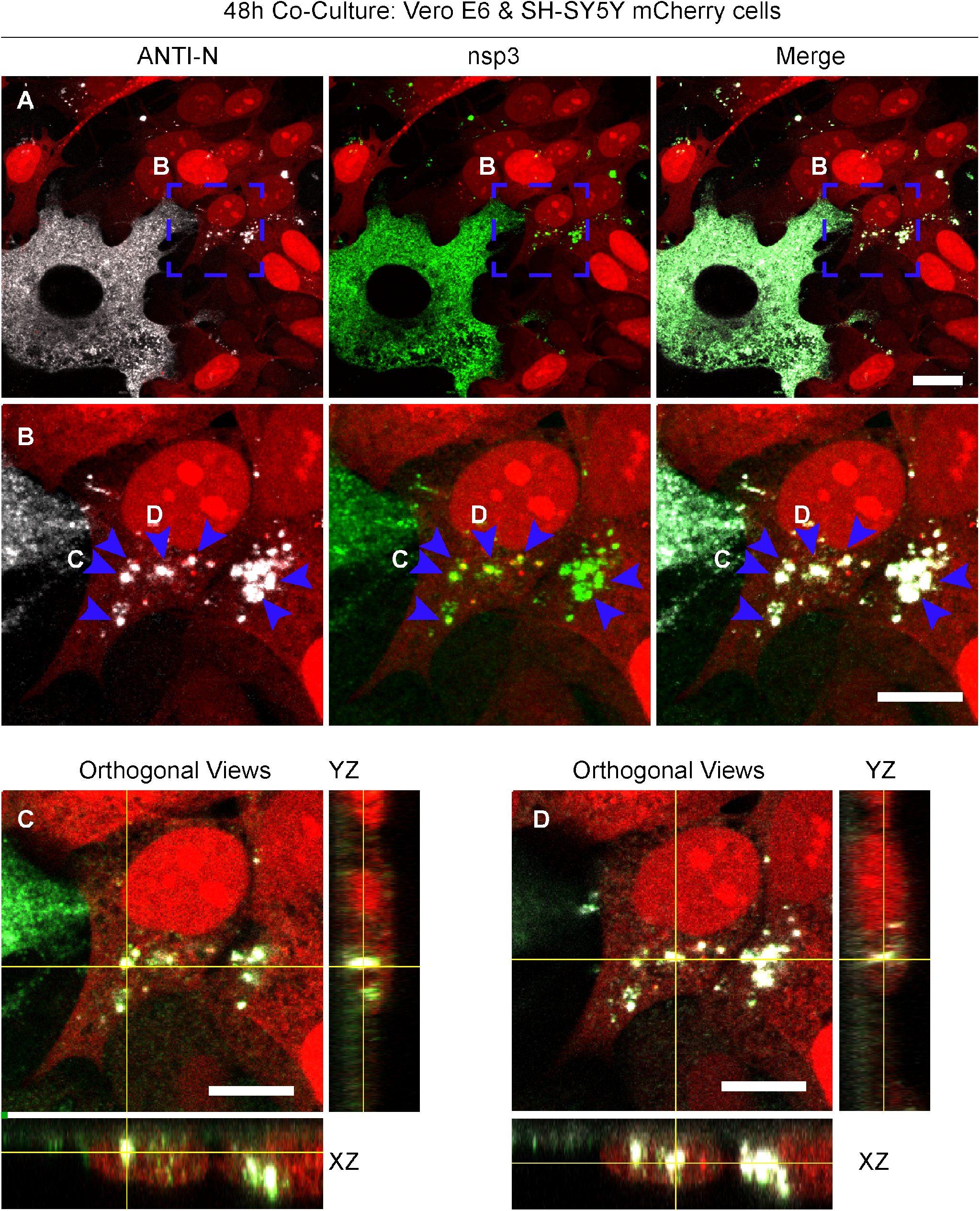
The non-structural protein 3 (nsp3) is detected in SH-SY5Y cells co-cultured with SARS-CoV-2 Vero E6 infected cells. (**A**) Confocal micrographs showing 48h co-culture of SARS-CoV-2 Vero E6 infected cells (donor) and SH-SY5Y mCherry cells (acceptor) stained by using anti-nsp3 and anti-N antibodies. (**B**) Enlargement of the blue dashed square in (A) showing puncta positive for both anti-nsp3 and anti-N in acceptor cells (**C-D**) Confocal micrographs representing the orthogonal views of (B) showing anti-nsp3 and anti-N puncta in the cytoplasm of acceptor cells. Blue arrowheads indicate anti-nsp3 and anti-N signal in acceptor cells. Scale bars, A, 10 µm, B-D, 10 µm.

### 2. TNTs contribute to the SARS-CoV-2 transmission

The results described above provide evidence that while SARS-CoV-2 entry into epithelial cells is mediated by the classical exo/endocytosis pathway, the spreading between permissive cells and non-permissive neuronal cells could occur in a direct cell-to-cell contact dependent manner. Recent studies have shown that viruses of many different families, including retroviruses such as HIV, and enveloped viruses like herpesviruses, orthomyxoviruses, and several others trigger the formation of TNTs or TNT-like structures in infected cells and use these structures to efficiently spread to uninfected cells (39). We therefore explored whether TNTs could be a mechanism involved in the spreading of SARS-CoV-2 to non-permissive neuronal cells. To detect TNTs between Vero E6 and SH-SY5Y cells we used cell Mask Blue, which stains the entire cell cytosol and outlines the entire cell shape. To properly identify TNTs by confocal microscopy it is crucial to distinguish them from other actin driven membranous protrusions such as filopodia (65, 66). TNTs hover above the substrate and even over other cells, and unlike dorsal filopodia, TNTs are able to connect two or more distant cells. Based on these criteria, first we assessed the presence of TNTs in control co-culture between Vero E6 cells and SH-SY5Y cells expressing the NLS-GFP (Nucleal Localization Signal) and stained with rhodamine phalloidin (for cellular membrane and actin labelling). We observed heterotypic TNTs formed between Vero E6 cells and SH-SY5Y NLS-GFP (fig. S5A, B). Strikingly, we also observed TNTs between SARS-CoV-2 infected Vero E6 donor cells and mCherry-SH-SY5Y acceptor cells in co-culture that contained particles stained with anti-N antibody, which were also found inside the cytoplasm of the acceptor neuronal cells (Fig. 4 A, B). In addition, we found TNTs between SH-SY5Y acceptor cells, which also contained anti-N labelled particles (Fig. 4 C, D). Altogether the data presented above (Fig 1-4) indicate that SARS-CoV-2 infection can be transferred in a cell-to-cell contact-dependent manner (likely TNT-mediated) from permissive cells to non-permissive neuronal cells. The evidence showing N-labelled particles in TNTs between neuronal cells suggests that once reached non-permissive cells infection could be further spread among them via TNTs.

**Fig. 4.**
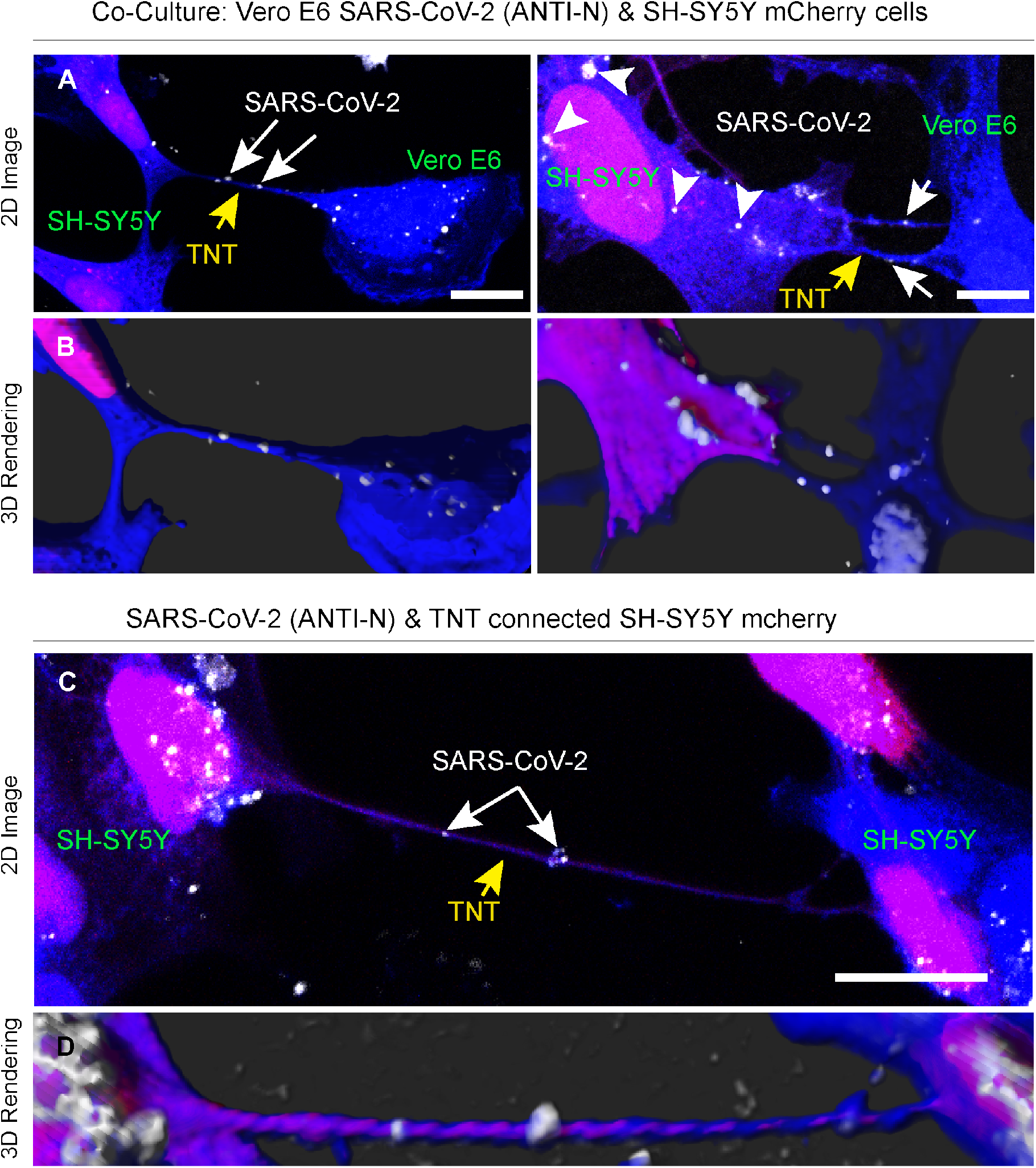
SARS-CoV-2 spread through TNTs from permissive infected Vero E6 to non-permissive SH-SY5Y mCherry cells. **(A, B)** SARS-CoV-2 infected Vero E6 cells (donor cells) were co-cultured with SH-SY5Y mCherry cells (acceptor cells). Co-culture were fixed at 24h (left top) and 48h (right top) and stained with the anti-N antibody to detect the virus. 2D confocal micrograph (A) and 3D rendering performed by using IMARIS software in (B) showing a TNT connecting SARS-CoV-2 infected Vero E6 cells and a SH-SY5Y mCherry cell; the yellow arrow points the TNT between Vero E6 and SH-SY5Y mCherry cells; the grey arrow indicates SARS-CoV-2 N signal inside TNT and in the acceptor cells. (**C, D)** 2D confocal micrograph (C) and 3D rendering performed by using IMARIS software (D) showing a TNT connecting two SH-SY5Y mCherry cells, co-cultured with Vero E6 infected cells. The yellow arrow points the TNT between the SH-SY5Y mCherry cells; the grey arrow indicates SARS-CoV-2 inside TNT. Scale bars A 10 µm, C, 15 µm.

### 3. Cryo-EM reveals viral compartments in TNTs between permissive and non-permissive cells

Next, we wanted to identify the nature and structure of the viral particles shared by TNTs and investigate the mechanisms allowing their TNT-mediated transfer to non-permissive cells (eg, whether the infectious particles were using TNTs as membrane bridges to surf on top, or as channels to be transferred inside the tube). To this aim we set up a correlative fluorescence and cryo-electron microscopy and tomography approach (CLEM, cryo-EM and cryo-ET) (35). These techniques allowed us to assess for the first time in correlative mode, both SARS-CoV-2 and TNTs architecture in the closest to native conditions. Vero E6 cells were infected with SARS-CoV-2 (MOI 0.05) and 48h post infection were seeded on Cryo-EM grids in co-culture with SH-SY5Y mCherry cells (Fig. 5A and 5 F). Before vitrification, the EM grids were imaged by fluorescence microscopy (FM) after labelling cells with either cell mask blue (Fig. 5A) or wheatgerm agglutinin (WGA) conjugated with a 488 fluorochrome (Fig. 5F) in order to identify TNTs (for details see material and methods). Consistent with our previous data, we detected TNTs between Vero E6 and SH-SY5Y mCherry (Fig. 5A, 5F). By using the grid finders after vitrification, we could identify precisely the TNTs position and image them at the ultrastructure level using both Glacios Cryo-TEM (Fig. 5A-E) and Titan Krios cryo-TEM (Fig. 5F-K). Strikingly, TNTs connecting Vero E6 infected cells and SH-SY5Y mCherry cells (labelled with cell mask blue) revealed the presence, inside the tube, of membranous structures of various sizes resembling DMVs (double-membrane vesicles) (Fig. 5D-E and Supplementary Movie 1). DMVs have been identified as the central hub for SARS-Cov2-RNA synthesis Klein et al, (58). Furthermore, as shown in the tomogram in figure 5I and in Supplementary Movie 2, the TNT also contained many vesicular structures (Fig. 5I-K blue arrow and Supplementary Movie 2). Of note, we never observed DMVs and such crowding of vesicular structures inside TNTs between Vero E6 and SH-SY5Y mCherry cells in control conditions (fig. S6 and Movie 3), where we could rather see isolated vesicles or organelles as in the case of the mitochondrion shown in fig. S6D and Movie 3.

**Fig. 5.**
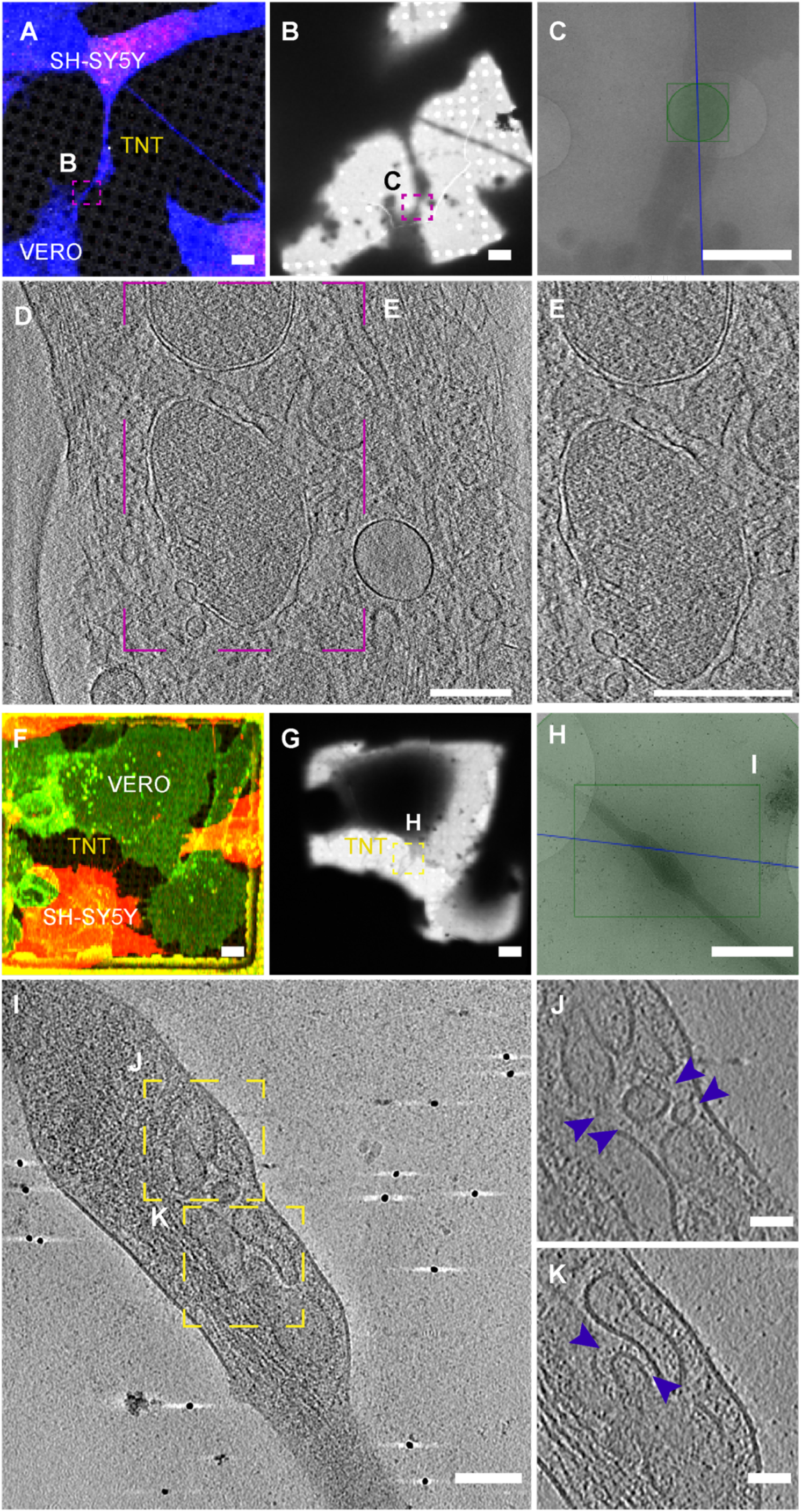
Ultrastructural analysis reveals SARS-CoV-2 viral compartments inside TNT between permissive Vero E6 cells and non-permissive SH-SY5Y neuronal cells. **(A)** Confocal micrographs showing a TNT connecting SARS-CoV-2 infected Vero E6 cells and SH-SY5Y mCherry cells stained with cell Mask blue. (**B)** Low and (**C)** intermediate magnification of an electron micrograph displaying TNT in (A). Green square in **(C)** corresponds to the high-magnification of cryo-tomogram slices in (**D)** showing a TNT containing vesicular compartments and double membrane vesicles (DMV). (**E)** Enlargement of cryo-tomogram slices in (D)**. (F-K)** Cryo-EM grids were prepared using Vero E6 infected cells co-cultured with SH-SY5Y mCherry cells and stained with WGA-488. (**F)** TNT between SARS-CoV2 infected Vero E6 cells and SH-SY5Y mCherry cells acquired by confocal microscopy (F) low (G) and intermedia (H) magnification TEM. (**I)** Slices of tomograms of TNT in the green square in (H) showing vesicular compartments inside TNT. (**J, K)** High-magnification cryo-tomography slices corresponding to the yellow dashed squares showing vesicular compartments inside TNT. The blue arrowheads indicate the vesicles inside TNT. Scale bars: A-C 2µm, D 200 nm, E 300 nm, F, G, H, 2µm, I 100 nm, J, K 50 nm.

Since SARS-CoV-2 replication is associated with proliferation of membranes and presence of DMVs where viral replication is taking place (58), it is possible that these structures represent viral replicative complexes being transferred to the acceptor cells.

### 4. TNTs facilitate SARS-CoV-2 transmission in permissive Vero E6 cells

Altogether, these data suggested that TNT can allow the intercellular spreading of the infection from permissive to non-permissive cells. Next, we wondered whether the TNT-mediated route dedicated to invading non-permissive cells could also be used to enhance the spreading of the virus between permissive cells in addition to the endocytic route. We therefore analysed whether SARS-CoV-2 exploits TNTs for cell-to-cell spread between Vero E6 cells (Fig. 6A-C). By confocal microscopy, we observed that in uninfected Vero E6 a low percentage of cells were connected by TNTs (Fig. 6A, C), but this percentage was substantially increased already after 24h of SARS-CoV-2 infection (Fig. 6B, C). Strikingly, after immunostaining using either the anti N antibody (Fig. 6B) and anti S (Fig. 6D) alone or both anti N and anti S antibodies (Fig. 6E) we could observe particles positive for one or both antibodies (Fig. 6B-E), suggesting that also mature virions could spread through TNTs between permissive cells. To verify the hypothesis that SARS-CoV-2, can use the TNT route to spread between permissive cells we decided to block the infection through the endocytic pathway. To this aim we used a neutralizing antibody which binds to the RBD domain of the spike protein (anti-SARS-CoV-2 human IgG C3 235) thus blocking binding to ACE2 receptor and the receptor-mediated entry of the virus. A preliminary experiment was set up to identify the minimal concentration of antibody sufficient to achieve neutralization of the viral stock of 1-5 x 10^5^ FFU/ml used to infect Vero E6 cells. The viral stock was incubated 1h at 37°C, 5% CO_2_ with three different concentrations of IgG C3 235 (1, 10 and 100 µg/ml) and then used to infect monolayers of Vero E6 cells for 48h. Viral production was then assessed by titration of the supernatant by focus forming assay (fig. S7) both 100 and 10 µg/ml concentration of antibody were enough to elicit complete neutralization of the viral stock, resulting in no sign of viral production (fig. S7). Therefore, a concentration of 10 µg/ml was set as the minimal concentration to investigate cell-to-cell transfer in Vero E6 cells.

**Fig. 6.**
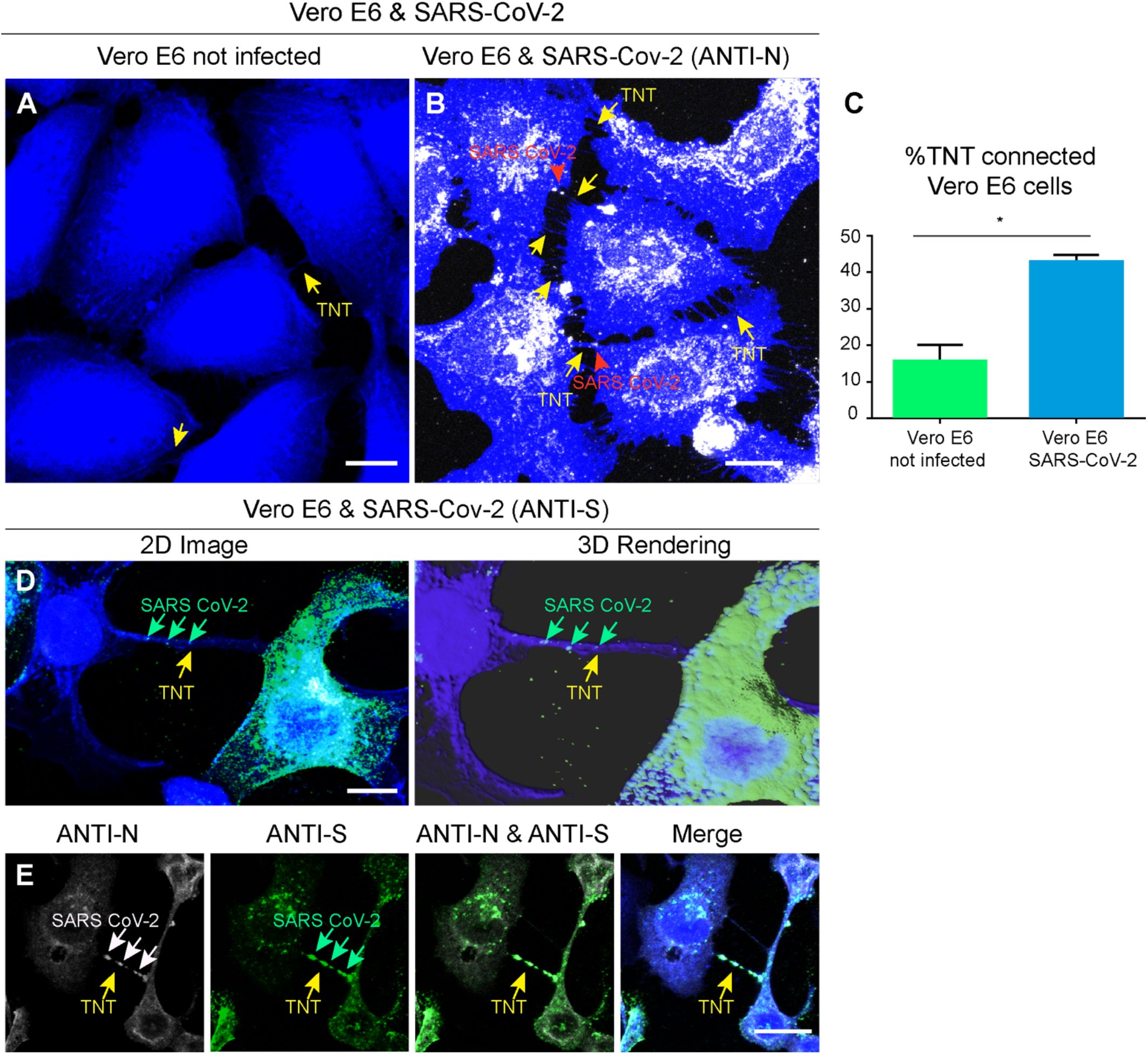
SARS-CoV-2 infection increases TNTs between infected Vero-E6 cells. **(A)** Confocal micrograph showing TNTs between non-infected Vero E6 cells. (**B)** Confocal micrograph showing TNTs between Vero E6 cells SARS-CoV-2 infected cells. Anti-N immunostaining is performed to detect SARS-CoV-2. The yellow arrows indicate TNTs between VeroE6 cells; the red arrowheads indicate SARS-CoV-2 signal associated to TNTs. (**C)** Graph showing the percentage of TNT-connected cells between Vero E6 cells non-infected and SARS-CoV2-infected. Mean percentage of TNT-connected Vero E6 non-infected cells: 13.95% ± 2.46. Mean percentage of TNT-connected SARS-CoV-2-infected Vero E6 cells: 44.69% ± 1.96 (***p=0.0006 for SARS-CoV-2-infected Vero E6 cells versus Vero E6 non-infected; N=3). (D) Confocal micrograph and 3D rendering showing TNTs between SARS-CoV-2-infected Vero E6 cells; an anti-S immunostaining was performed to detect SARS-CoV-2. The yellow arrow indicates a TNT between Vero E6 infected cells; the green arrows indicate SARS-CoV-2 associated to a TNT. (**E)** Confocal micrograph showing TNTs between Vero E6 cells SARS-CoV-2 infected cells labelled with cell Mask blue. Anti-N (633) and anti-S (488) immunostaining was performed to detect SARS-CoV-2. The yellow arrows indicate a TNT between infected VeroE6 cells; the white and the green arrows indicate SARS-CoV2 particles inside TNTs. Scale bars A, B, E 15 µm, C 10 µm.

As usual, Vero E6 cells were infected with SARS-CoV-2 MOI 0,05 (donor cells) for 48 hours, but then incubated with 10 µg/ml anti-SARS-CoV-2 IgG C3 235 for 1 hour at 37°C 5% CO_2_, prior co-culturing in presence of the antibody, with Vero E6 cells expressing mCherry (acceptor cells), to distinguish them from the infected donor population. After 24h and 48h hours cells were fixed and immunostained with anti-N antibody. By using confocal microscopy and ICY software we evaluated the percentage of Vero E6 acceptor cells containing anti N positive viral particles both in control condition and in the presence of the blocking antibody (Fig. 7A-C). Interestingly, after 24h of co-culture in presence of the anti-S neutralizing antibody, 42,9% of acceptor cells were positive for SARS-CoV-2 detected by anti-N immunostaining, and this percentage increased to 63,8% after 48h of co-culture, compared to co-culture control conditions (not incubated with the anti-S neutralizing antibody) where respectively 95% at 24h and 96,8% at 48h of acceptor cells were positive for anti-N immunostaining (Fig. 7C). As control, to verify the absence of infectious virus in the supernatant of the coculture treated with the anti-S neutralizing antibody, we challenged naïve Vero E6 cells with the supernatants of both the co-cultures (incubated/and not with the 235 Ab) (Fig. 7D-F). While 100% of cells infected with the untreated supernatant were positive for SARS-CoV-2 (Fig. 7D, F), no infection could be detected in the cells challenged with the treated supernatant (Fig. 7E, F). Furthermore, the supernatants of each condition were collected to determine the virus concentration using the focus forming assay (Fig. 7 G).

**Fig. 7.**
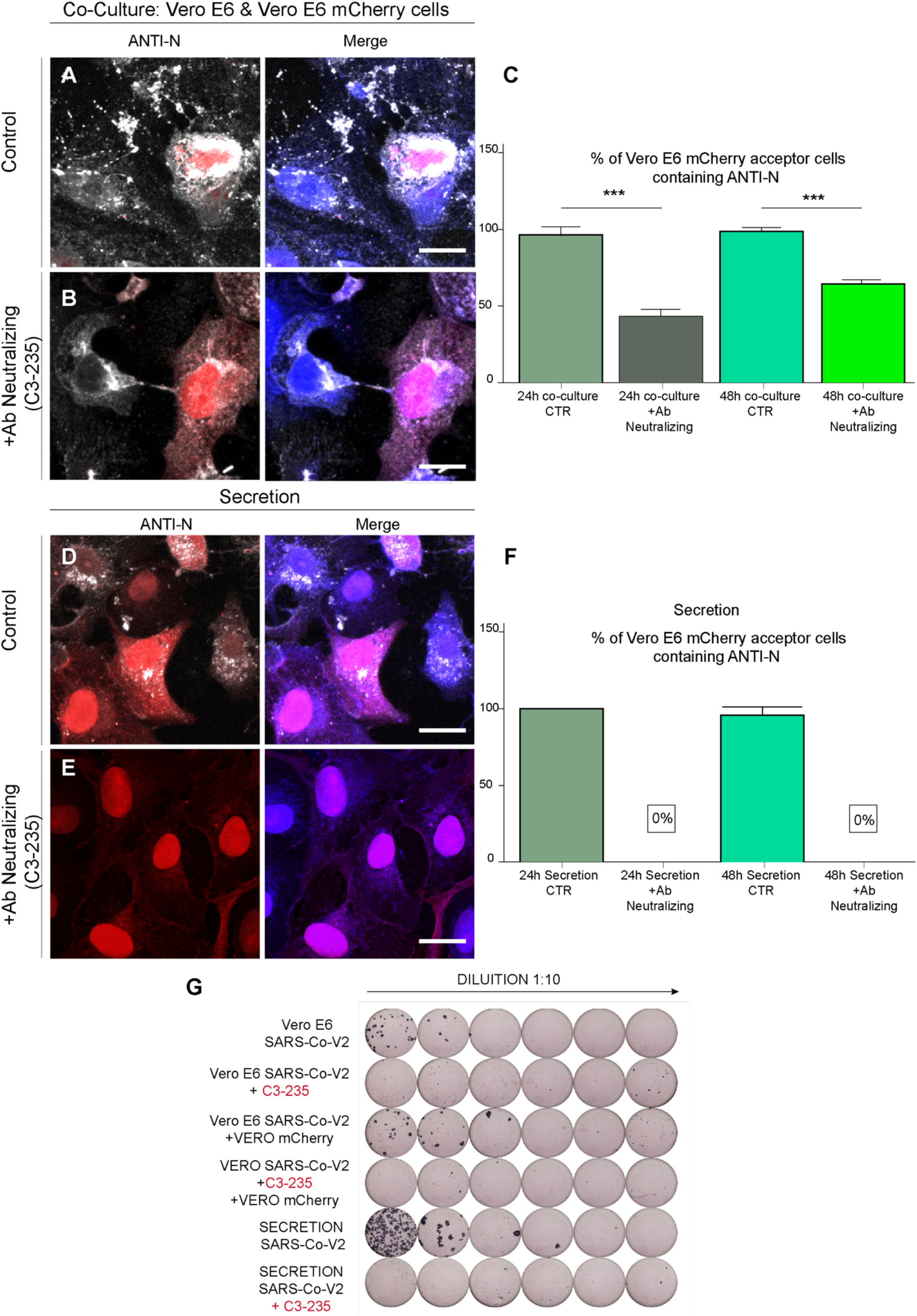
SARS-CoV-2 spread through TNTs between permissive cells. **(A)** Vero E6 infected donor cells were put in co-culture at 1:1 ratio with Vero E6 mCherry acceptors in control conditions (without neutralizing antibody) and (B) in neutralizing conditions. (**B**) Donor Vero-E6 infected cells were incubated with 10 µg/ml of anti-SARS-CoV-2 IgG C3 235 for 1 hour at 37°C 5% CO2 before to be co-cultured with Vero E6 mCherry acceptors cells. The co-cultures were fixed after 48h of incubation at 37°C 5% CO2 and immunostained with anti-N antibody to detect SARS-CoV-2. (**C**) Graph showing the mean percentage of N puncta transferred in co-culture at 24h and 48h, treated and not with the neutralizing antibody. 24h co-culture control: 95.45% ± 4.29 versus 24h co-culture plus neutralizing antibody: 42.91 ± 4.55 (**p=0.0018 24h co-culture control versus 24h co-culture plus neutralizing antibody). 48h co-culture control: 96.88% ± 3.12 versus 48h co-culture plus neutralizing antibody: 63.90 ± 1.99 (*p=0.0104 48h co-culture control versus 48h co-culture plus neutralizing antibody. (D, E) Secretion test; (**D**) Vero E6 mCherry cells were incubated with the supernatant deriving from donor Vero E6 infected cells. (**E**) The supernatant from donor Vero E6 infected cells was incubated with 10 ug/ml of the anti-SARS-CoV-2 IgG C3 235 for 1h at 37°C, in order to neutralize the viral particles, before to be added on top of Vero E6 mCherry acceptor cells. After 48h of incubation at 37°C 5% CO2, the secretion samples were fixed and immunostained for anti-N. (**F**) Graph showing the mean percentage of N puncta contained in acceptor cells in the secretion experiments at 24h and 48h, treated or not with the neutralizing antibody. 24h control: 100% ± 0 versus 24h plus neutralizing antibody: 0% ± 0 (***p=0.0005 24h control versus 24h plus neutralizing antibody). 48h control: 80% ± 10 versus 48h plus neutralizing antibody: 0% ± 0 (***p=0.0008 48h control versus 48h plus neutralizing antibody). (**G**) Supernatant of each condition was then collected to assess viral neutralization using Focus Forming Assay (FFA) titration protocol. Scale bars A-E 20 µm.

These data show that despite the block of the receptor-mediated virus entry, Vero E6 cells could be infected when put in co-culture with infected cells, indicating that SARS-CoV-2 can spread also between permissive cells through a secretion-independent pathway.

### 5. Cryo-EM reveals SARS-Cov-2 on top of TNTs

To uncover the nature of the particles labelled with anti-N and anti-S antibodies in TNTs by fluorescence microscopy and to discriminate if they were localised inside or on top of TNTs between permissive cells we used again correlative fluorescence cryo-EM and cryo-ET (35). Vero E6 cells were infected with SARS-CoV-2 (MOI 0.05) and 48h post infection seeded on Cryo-EM grids. After having identified by FM the exact location of the TNT connecting Vero E6 infected cells (Fig. 8A), the EM-grids were cryo-fixated and analysed by cryo-EM. High-quality 3D images using a Titan Krios Microscope revealed SARS-CoV-2 viral particles located on the surface of TNTs-connecting two Vero E6 cells (Fig. 8 D-H, fig. S8, Movie 4 and 6). SARS-CoV-2 particles that decorate the TNTs’ surface displayed both an ellipsoidal and spherical enveloped morphology with an average diameter ranging from 50 to 100 nm typically of a coronavirus (Fig. 8, fig. S8, Movie 4 and 6). In our tomograms and in the Supplementary Movie 4, 6 we can clearly discern the most distinctive features of the virus: the spike proteins that decorate the surface of the viral particles, together with the ribonucleoprotein complexes (RNPs) organized inside the virus (Fig. 8 D-H, fig. S8, Supplementary movie 4 and 6) in accordance with recent cryo-EM data for the virus isolated from infected cells (67–69) and cryo-FIB SEM pictures of the intracellular virus (58).

**Fig. 8.**
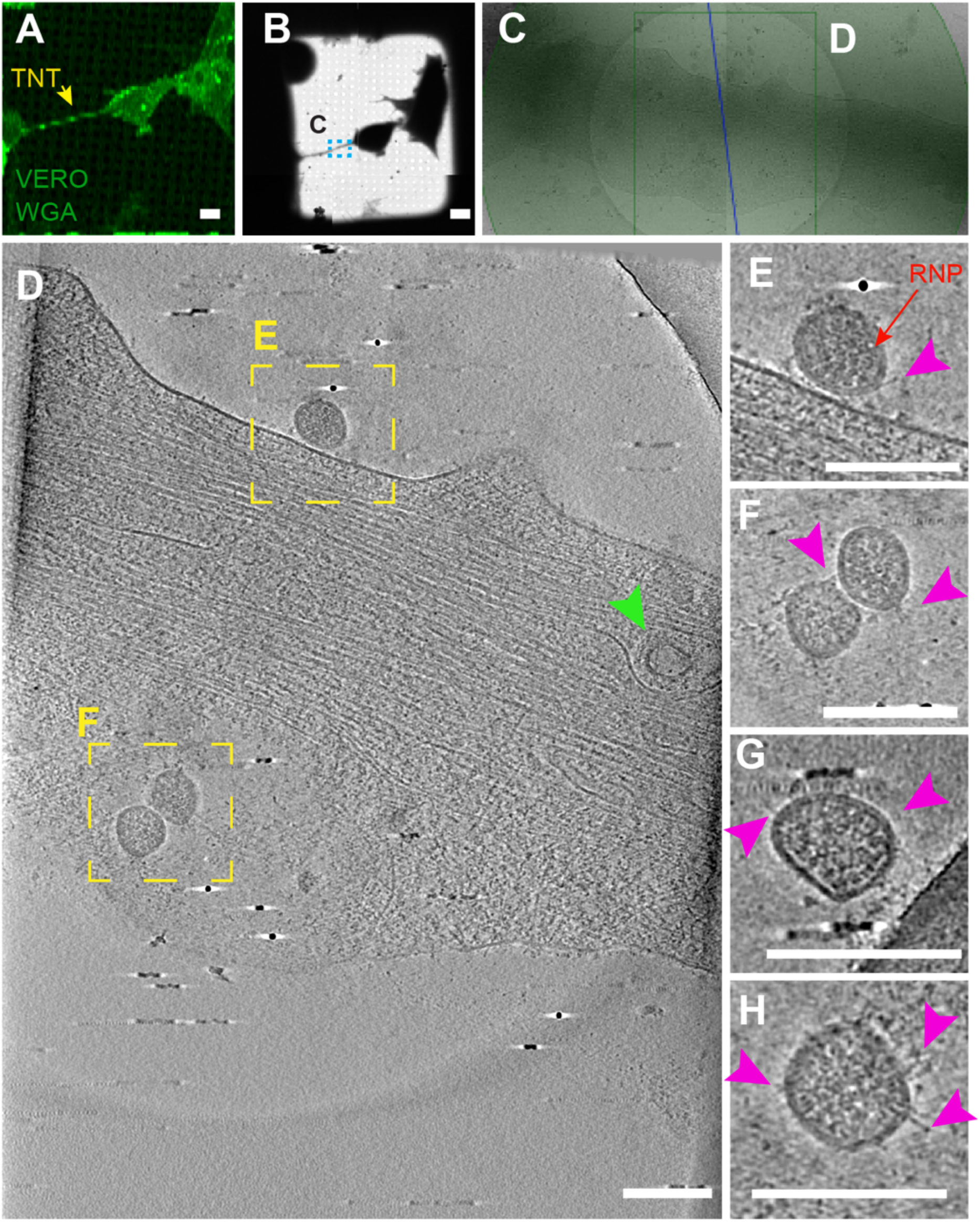
Correlated light and cryo-electron microscopy strategies reveal SARS-CoV-2 on top of TNTs between infected Vero E6 cells. **(A)** TNT-connected Vero E6 infected SARS-CoV-2 cells stained with WGA (green) and acquired by confocal microscopy (**A**) low (**B**) and intermediate (**C**) magnification TEM. (**D**) Slices of tomograms of TNT in green square in (**C**) showing extracellular SARS-CoV-2 on top of the TNT connecting Vero E6 cells. (**E-F)** High-magnification cryo-tomography slices corresponding to the yellow dashed squares in (**D)** showing SARS-CoV-2; RNP proteins and Spike are observed. Pink arrowheads indicate the spike; red arrow points the RNP proteins. **(E, F, G, H)** High-magnification cryo-tomography slices showing the extracellular virions on TNT. Scale bars, A, B, J, K 10 µm, C, I 2µm, D-H 150 nm, L 200 nm, M 50 nm.

The presence of the virus on top of TNTs was observed only in permissive cells, and not in neuronal cells (Fig. 5). This difference could be explained by the presence of the ACE2 receptor, which is only expressed on the cell surface and TNT membranes of Vero E6 cells and not on SH-SY5Y cells (fig. S2). We also observed vesicular structures (average diameter of 50-100 nm) inside TNTs connecting Vero E6 infected cells (Fig. 8D green arrowhead and Movie 4), similar to the ones observed in the TNT between Vero E6 infected cells and SH-SY5Y mCherry cells (Fig. 5D-F and Movie 2). As the observation inside TNTs is more challenging compared to the analysis of the TNT surface, in order to unequivocally demonstrate that these vesicular structures corresponded to the virus and/or viral compartments, we set-up a challenging correlative IF cryo-EM protocol making use of the anti-S antibody. SARS-CoV-2-infected Vero E6 cells, seeded on EM grids, were fixed and processed for an immunostaining against the Spike proteins (anti-S). Before cryo-fixing the EM grids, we selected the TNTs connecting cells that were positive for the anti-S by using confocal microscopy (Fig. 9A) and then observed in the same position at cryo-EM (Fig. 9B-D). In correspondence of the fluorescent anti-S antibody signal in TNTs coming from a Vero E6 cells (Fig 9A), we could observe structures resembling SARS-CoV-2 particles that decorate the TNTs’ surface with both a spherical and an ellipsoidal enveloped morphology and spike-like structures (Fig. 9D-I and Movie 5). Note that the images here are less clear compared to figure 8 and supplementary figure 8 as in this case the grids were acquired with Glacios Cryo-TEM instead of Titan Krios cryo-TEM. Interestingly, in correspondence of the S antibody we observed multiple vesicular structures inside the TNT (Fig. 9D, H-L and Movie 5), similar to the ones observed inside TNTs between permissive and non permissive cells (Fig. 5I-K blue arrow and Supplementary Movie 2). Considering that cryo-ET resolution is limited by the thickness of the sample, and the fact that TNTs described here have an average diameter of more than 500 nm, we were at the resolution limits and we were not able to discriminate the precise structures of these vesicular compartments. Therefore, besides stating the fact that these structures were found in correspondence of S antibody, we cannot be sure whether they are mature virions as the ones observed outside the TNTs.

**Fig. 9.**
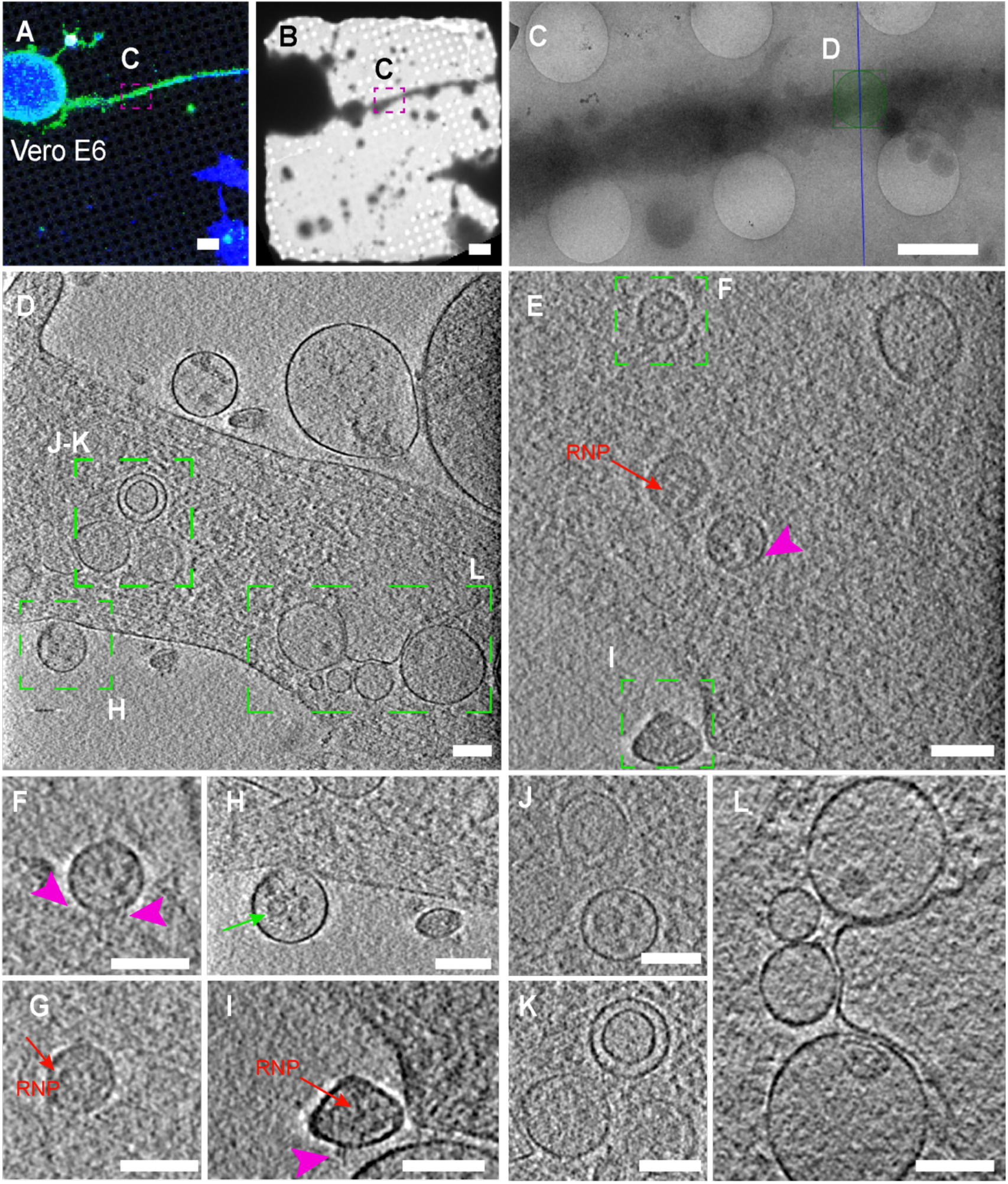
Correlative IF cryo-EM strategies to discriminate SARS-CoV-2 localization in TNTs. (**A-L** Cryo-EM grids were prepared using Vero E6 infected cells (**A)** Confocal micrograph showing TNT connecting infected Vero E6 cells stained with anti-S antibody (green) and cell mask blue. Low (**B**) and intermediate (**C**) magnification TEM of (A). (**D-E)** High-magnification cryo-tomography slices showing vesicular compartments in correspondence of anti-S signal and SARS-CoV-2 like structure on TNT surface. (**F-I)** Enlargement of the high-magnification cryo-tomography slices (D-E) showing SARS-CoV-2 like structure on TNT surface (**J-L)** Enlargement of the high-magnification cryo-tomography slices (D-E) showing viral like structure inside TNT. RNP proteins and Spike are observed. Pink arrowheads indicate the spike; red arrow points the RNP proteins, green arrow indicates SARS-CoV-2 like structure inside vesicles. Scale bars, A, B 10 µm, C, 2µm, D-L 100 nm.

Our observations are in line with the recent publication by Caldas and colleagues showing by classical scanning electron microscopy that SARS-CoV-2 particles appear surf on top of cellular protrusions (70). However, in this case the authors did not enquire whether these protrusions were TNTs and whether they allowed viral transfer. Thus, despite several reports describing virus of different families (71, 72) on top of filopodia and/or cellular extension, including SARS-CoV-2 (58,70,73), our report represents the first evidence that TNTs could be one possible route for the spreading of SARS-CoV-2.

Our study is carried out using Vero E6 cell line as epithelial model since it has been widely employed for SARS-CoV-2 isolation, propagation, and antiviral testing, due to its high virus production and a prominent cytopathic effect (CPE) upon infection (46). Our choice of SH-SY5Y cell line as neuronal model of non-permissive cells is justified not only because these cells are human and widely used as neuronal model, but also because we have thoroughly characterized their TNTs (35) and able to identify them with high reliability (35, 74). An ideal neuronal model could be represented by primary neurons, however in primary neurons it is very difficult to discriminate TNT-like structures (32) and it is even more challenging to apply correlative cryo-light and electron microscopy and–tomography approach. In the future it will be important to confirm these data using a different neuronal model like iPSC-derived human neurons that are able to recapitulate the complexity of human neurons. Furthermore, as we demonstrated here that TNTs allow the passage of the virus between permissive cells this work sets the basis to investigate the potential role of TNTs in allowing the spreading of the virus from the olfactory epithelium of the nasal cavity to the olfactory sensory neurons in the central nervous system (CNS).

Overall, within the limitation of our study mentioned above, our report provides unique structural information of SARS-CoV-2 and how it could use TNTs for its spreading between permissive cells and non-permissive cells (e.g., neuronal cells), thus increasing both viral tropism and infection efficiency. These results also pave the way to further investigations on the role of cell-to-cell communication in SARS-CoV-2 spreading to the brain in more physiological contexts, and on alternative therapeutically approaches to impair viral spreading in addition to the current investigations mainly focused on blocking the spike-receptor interactions.

## Methods

### Cell Lines and Viruses

African green monkey kidney Vero E6 cell and Colorectal Adenocarcinoma human epithelial (Caco-2) cells were maintained at 37 °C in 5% CO2 in Dulbecco’s minimal essential medium (DMEM) (Sigma) supplemented with 10% fetal bovine serum (FBS) and 1% penicillin/streptomycin.

Human neuroblastoma (SH-SY5Y) cells were cultured at 37 °C in 5% CO2 in RPMI-1640 (Euroclone), plus 10% fetal bovine serum and 1% penicillin/streptomycin. Mouse catecholaminergic neuronal cell line, Cath.a-differentiated cells (CAD) were kindly given by Hubert Laude (Institut National de la Recherche Agronomique, Jouy-en-Josas, France) and cultured at 37 °C in 5% CO2 in Gibco Opti-MEM (Invitrogen), plus 10% fetal bovine serum and 1% penicillin/streptomycin.

The strain BetaCoV/France/IDF0372/2020 was supplied by the National Reference Centre for Respiratory Viruses hosted by Institut Pasteur (Paris, France) and headed by Pr. Sylvie van der Werf. The human sample from which strain BetaCoV/France/IDF0372/2020 was isolated has been provided by Dr. X. Lescure and Pr. Y. Yazdanpanah from the Bichat Hospital, Paris, France. Moreover, the strain BetaCoV/France/IDF0372/2020 was supplied through the European Virus Archive goes Global (Evag) platform, a project that has received funding from the European Union’s Horizon 2020 research and innovation programme under grant agreement No 653316.

### Viral infection to identify SARS-CoV-2 permissive cells

In order to assess which cell-lines were permissive to SARS-CoV-2 infection, the different cells were plated on a 96 multiwell plate and infected with a MOI (multiplicity of infection) from 10^-1^ to 10^-5^ in DMEM 2% FBS. The cell-lines used in this assay included Caco-2, CAD, SH-SY5Y and Vero E6. All the cells were plated at a 60% confluence. The cells were incubated in infection media for 3 days. At day 2 and 3 post infection an aliquot of the supernatant from the higher MOI was collected for titration. At day 3 p.i. the monolayers were then fixed with 4% PFA and viral infection was visualized using an immunofluorescence protocol.

### Immunofluorescence protocol for Immunospot

After 45 minutes incubation with 4% PFA, the monolayers were washed with PBS and incubated 5 minutes with PBS 1X-0.5% Triton at R.T. (Room Temperature); the cells were then washed and incubated for 10 minutes with PBS 1X-50mM NH4Cl. After washing, 30 minutes blocking was performed using PBS 1X-2% BSA; the monolayers were incubated with the primary antibody, a polyclonal SARS-CoV-antiN IgG, provided by Nicolas Escriou, Institut Pasteur, Paris, overnight at 4 degrees. After washing, the cells were then incubated with a goat anti-rabbit Alexafluor488-conjugated antibody for 1 hour. After washing with PBS 1X to remove the unbound antibody, the immunofluorescence was visualized using the Fluoro-X suite of a C.T.L. Immunospot® S6 Image Analyzer.

### Semi-solid Plaque Assay

The aliquots of supernatant collected at day 2 and day 3 were used to assess viral production through a semisolid plaque assay. Each sample underwent 1:10 serial dilutions. 250 μl of each dilution was used to infect a confluent monolayer of Vero E6 cells, in a 24 wells multiwell plate, for a total of 6 wells per sample.

Viral absorption was allowed for 1 hour at 37°C, and a semisolid overlay, composed by MEM 1X, 10% FBS and 0.8% agarose, was then added to the infection (250 μl per well). The cells were incubated at 37°C 5% CO2 for 72 hours. Finally, the infected monolayers were fixed with 500 μl of 4 % PFA for 30 minutes. Afterward, the PFA was removed and the monolayers were then stained with Crystal Violet solution containing 2 % PFA, to evaluate the cytopathic effect. The reaction was stopped after 15 mins and residual Crystal violet was removed through immersion in diluted bleach, followed by washing in water.

### Focus forming assay

Vero E6 cells were plated in a 96 multiwell plate 2 x 10^4^ cells per well. The monolayers were then infected with serial dilutions (1:10) of samples to be titrated. The infection was allowed for two hours at 37°C 5% CO2. Afterwards, the infection medium was removed and a semisolid overlay composed by 1.5% Carboxymethyl-cellulose and MEM 1X was added to the monolayer. The cells were incubated for 36 hours at 37°C 5% CO2 to allow foci formation. The monolayers were then fixed with 4% paraformaldehyde; after 45 minutes they were washed with PBS and incubated 5 minutes with PBS 1X-0.5% Triton at R.T. (Room Temperature); the cells were then washed again and incubated for 10 minutes with PBS 1X-50mM NH4Cl. After washing, the cells were incubated 2 minutes in 0.05% PBS-tween and then incubated with the primary antibody, a polyclonal SARS-CoV-anti-N IgG, provided by Nicolas Escriou, Institut Pasteur, Paris (or alternatively with a Human SARS-CoV-2 anti-S IgG provided by Cyril Planchais from the group of Hugo Mouquet Institut Pasteur, Paris), overnight at 4 degrees. After washing the cells were then incubated with an anti-rabbit (or an anti-human) HRP-conjugated antibody for 1 hour. After washing with PBS 1X to remove the unbound antibody, the foci were visualized using a DAB staining solution in PBS with 8% NiCl, and washed 3 times with water to stop the reaction. The foci were then visualized and counted using the Biospot suite of a C.T.L. Immunospot® S6 Image Analyzer.

### Lentiviral Transduction

Transduction of SH-SY5Y and Vero E6 cells with a lentiviral vector expressing pCMV-mcherry: 600.000 SH-SY5Y cells 400.000 Vero-E6 were plated in 60 mm plates. After 24 h, they were infected with 800 µl of LV-pCMV-mCherry. After 48 h, cells expressing mCherry have been validated. Transduction of SH-SY5Y cells with a lentiviral vector expressing pCMV-H2B-GFP: 600.000 SH-SY5Y cells were plated in 60 mm plates. After 24 h, they were infected with 800 µl of LV-pCMV-H2B-GFP. Transduction of SH-SY5Y cells with a lentiviral vector expressing pCMV-H2B-GFP: 600.000 SH-SY5Y cells were plated in 60 mm plates. After 24 h, they were infected with 800 µl of LV-pCMV-H2B-GFP.

### SARS-CoV-2 infection of Vero E6 cells for co-culture experiments and cryo-EM grids

1.000.0000 of donor Vero E6 cells were infected with a MOI of 0.05 in DMEM 1% FBS for 2 hours. Afterward, the infection medium was removed and substituted with fresh DMEM 10% FBS. The cells were left in incubation at 37°C with 5% CO2 for 48 hours. After that time, cells were trypsinzed, centrifuged (1000 rpm, 10 min), counted and seeded for the different experiments.

### Co-culture preparation for SARS-CoV-2 transfer experiments and Secretion test

1.000.0000 of donor Vero E6 cells were infected with a MOI of 0.05 in DMEM 1% FBS for 2 hours. Afterward, the infection medium was removed and substituted with fresh DMEM 10% FBS. The cells were left in incubation at 37°C with 5% CO2 for 48 hours. As acceptors were used the non-permissive SH-SY5Y cells and permissive Vero-E6 cells stably transfected with a lentivirus expressed mCherry, according to the kind of experiment. The infected donors, as well as the acceptors cells, were trypsinized centrifuged (1000 rpm, 10 min), counted and co-cultured on 24 glass coverslips 37°C with 5% CO2with 1:1 ratio (50.000 donor-50.000 acceptor). After 24h and 48h, co-cultures were washed with 0.01% trypsin to remove excess of virus on top of the cell membrane and fixed for 30 min with 4% PFA, then we proceed processing the co-culture for an immunostaining for anti-Nucleoprotein and anti-Spike. After the immunostaining, cells were stained with HCS Cell Mask TM Blue Stain (Invitrogen, 1:300) in PBS1X for 30 min then 30 mounted.

Images were acquired on an LSM 700 confocal microscope (Zeiss) with a 40X objective. After image acquisition, number of acceptor cells, which had received SARS-CoV-2 identified by the anti-N and/or anti-S immunostaining were quantified. Automated detection and quantification of the number of acceptors received SARS-CoV-2 was assessed with the open source software, ICY as described above.

To evaluate the possibility of SARS-CoV-2 transfer from donor to acceptor cells mediated by secretion, the supernatants from SARS-CoV-2 infected Vero-E6 cells were collected centrifuged at 1000 rpm for 10 min to remove floating cells and added on acceptor cells: SH-SY5Y mCherry. After 24h and 48h acceptor cells, acceptor cells were washed with 0.01% trypsin and fixed with 4% PFA at RT for 30 min. After image acquisition, acceptor cells were counted for the presence of SARS-CoV-2 signal. Secretion test was performed in parallel to all the co-coculture experiments performed in this study by following the same protocol.

Additionally, the supernatants from donor infected cells were used to assess viral production by focus forming assay titration protocol.

### Immunofluorescence labeling

Cells were fixed in 4% PFA for 30 min, quenched with 50 mM NH4Cl for 15 min, permeabilized with 0,5% Triton-100 for 5 min in PBS 1X, blocked with PBS 1X containing 2% BSA (w/v) for 1 h. Cells were then incubated with primary antibody dissolved in 2% BSA in PBS-1X. The primary antibody used were: a rabbit anti-Nucleoprotein (Anti-N, gift from Nicolas Escriou, Institut Pasteur, Paris) (1:500) over night (ON); an anti-human anti-Spike (H2-162, produced by Cyril Planchais from the group of Hugo Mouquet Institut Pasteur, Paris) (1: 100) ON, an anti-mouse J2 (1:50) (Scicons) ON, an anti-sheep nsp3 (1:200) (MRC PPU Reagents).

The day after, cells were thoroughly washed and incubated for 40 min with an anti-rabbit Alexa-Fluor 633-conjugated secondary antibody (Invitrogen), an anti-human Alexa-Fluor 488-conjugated secondary antibody (Invitrogen), goat anti-mouse Alexa-Fluor 633-conjugated secondary antibody (Invitrogen) at 1:500 in 2% BSA (w/v) in PBS 1X respectively. Cells were then carefully washed in PBS 1X and labeled with HCS Cell Mask TM Blue Stain (Invitrogen, 1:300) in PBS 1X for 30 min then 30 mounted. For ACE2 Antibody (PA5-20046-Thermo Fisher Scientific) Immunostaining: cells were fixed in 4% PFA for 10 min, quenched with 50 mM NH4Cl for 15 min, blocked with PBS 1X containing 2% BSA (w/v) for 1 h. Cells were then incubated with primary antibody ON dissolved in 2% BSA in PBS-1X. The day after, cells were thoroughly washed and incubated for 40 min with an anti-rabbit Alexa-Fluor 488-conjugated secondary antibody (Invitrogen). Cells were then carefully washed in PBS 1X and labeled with HCS Cell Mask TM Blue Stain (Invitrogen, 1:300) in PBS 1X for 30 min then 30 mounted.

### Co-culture preparation for SARS-CoV-2 transfer experiments in presence of neutralizing antibody

Vero-E6 donor cells, infected as previously described, were put in co-culture, in a 1:1 ratio, with mCherry-Vero-E6 acceptors in presence of a SARS-CoV-2 neutralizing antibody. Briefly, infected donors were trypsinized and counted. They were then diluted at a concentration of 5 x 10^5^ cells per ml in DMEM 5% FBS, containing a concentration of 10 ug/ml of anti-SARS-CoV-2 IgG C3 235 [produced by Cyril Planchais from the group of Hugo Mouquet Institut Pasteur, Paris] which it has been proved to be sufficient to elicit complete neutralization for a viral concentration of 1-5 x 10^5^ FFU/ml. Donor cells were incubated in presence of the antibody for 1 hour at 37°C 5% CO2. Afterward, donor cells were co-cultured in a ratio 1:1 with mCherry-Vero-E6 acceptor cells in DMEM 5% FBS with 10 ug/ml of the afore mentioned neutralizing antibody. The co-cultures were incubated for 24 and 48h hours 37°C 5% CO2. Then, co-cultures were fixed in 4% PFA for 30 min and immunostained for the anti-N (protocol described above) and with HCS Cell Mask TM Blue Stain (Invitrogen, 1:300). Images were acquired on an LSM 700 confocal microscope (Zeiss) with a 40X objective. After image acquisition, number of acceptor cells, which had received SARS-CoV-2 identified by the anti-N immunostaining were quantified ICY software as before. In parallel, supernatant of each conditions was then collected to assess viral neutralization using Focus Forming Assay (FFA) titration protocol. For the secretion test, performed in parallel with the co-culture, an aliquot of the supernatant from donor was incubated with 10 ug/ml of the anti-SARS-CoV-2 IgG C3 235 for 1h at 37°C, in order to neutralize the viral particles, present in the supernatant. In parallel another aliquot was left untreated, for comparison. The supernatants were then added on top of acceptors cells. Afterwards, we proceed for the analysis as before mentioned.

### TNT counting

For quantification of TNT-connected cells, Vero-E6 cells infected (as described before) and not infected were trypsinised and counted; 50,000 cells were plated on 24 glass coverslips. After 24h, cells were fixed (15 min at 37 °C in 2% PFA, 0.05% glutaraldehyde and 0.2 M HEPES in PBS, and then additionally fixed for 15 min in 4% PFA and 0.2 M HEPES in PBS). Cells were carefully washed in PBS, labeled for 20 min at RT with a 3.3 µg/µL solution of Wheat-Germ Agglutinin (WGA) Alexa Fluor©-647 nm conjugate (Invitrogen) in PBS, washed again and mounted. The whole cellular volume was imaged by acquiring 0.45 µm Z-stacks with an inverted confocal microscope (Zeiss LSM 700) using ZEN software. TNT-connected cells, cells connected by straight WGA-labeled structures that do not touch the substrate, were manually counted by ICY software using semi-automatized TNT counting tool as previously described (36, 65). The 3D rendering of TNTs were performed using IMARIS software.

### Cell preparation for cryo-EM

Carbon-coated gold TEM grids (Quantifoil NH2A R2/2) were glow-discharged at 2 mA and 1.5–1.8 × 10-1 m bar for 1 min in an ELMO (Cordouan) glow discharge system. Grids were sterilized under UV three times for 30 min at RT and then incubated at 37 °C in complete culture medium for 2h. 200,000 Vero-E6-infected cells (after 48h post infection) were counted and seed on cryo-EM grids positioned in 35 mm Ibidi µ-Dish (Biovalley, France). For co-culture, 100,000 Vero E6-infected cells (after 48h post infection) were co-cultured with 100,000 mCherry-SH-SY5Y on cryo-EM grids in 35 mm Ibidi µ-Dish (Biovalley, France). After 24h of cells resulted in 3 to 4 cells per grid square. Prior to chemical and cryo-plunging freezing, cells were labeled with WGA-Alexa-488 (1:300 in PBS) for 5 min at 37 °C. For correlative light-and cryo-electron microscopy, cells were chemically fixed in 2% PFA + 0.05% GA in 0.2 M Hepes for 15 min followed by fixation in 4% PFA in 0.2 M Hepes for 15 min and kept hydrated in PBS buffer prior to vitrification.

For correlative light-and cryo-electron microscopy using the anti-S primary antibody, cells were fixed with PFA 4% for 15 min at 37 °C, quenched with 50 mM NH4Cl for 15 min, and blocked with PBS containing 2% BSA (w/v) for ON at 4 °C. Cells were labeled with an anti-human AlexaFluor 488-conjugated secondary antibody (Invitrogen) at 1:500 and labelled with HCS Cell Mask TM Blue Stain (Invitrogen, 1:300). For cell vitrification, cells were blotted from the back side of the grid for 8 s and rapidly frozen in liquid ethane using a Leica EMGP system as we performed before (35).

### Cryo-electron tomography data acquisition and tomogram reconstruction

The cryo-EM data was collected from different grids at the Nanoimaging core facility of the Institut Pasteur using a Thermo Scientific Titan Krios G3i electron microscope with a Gatan Bioquantum energy filter and K3 detector. Tomography software from Thermo Scientific was used to acquire the data. Tomograms were acquired using dose-symmetric tilt scheme (75), a +/-60 degree tilt range with a tilt step 2 was used to acquire the tilt series. Tilt images were acquired in counting mode with a calibrated physical pixel size of 3.2 Å and total dose over the full tilt series of 3.295 e- /Å2 and dose rate of 39,739 e-/px/s with an exposure time of 1s. The defocus applied was in a range of −3 to - 6 µm defocus.

The tomograms showed in the Figure 8 and 9 were performed on Glacios equipped with a field emission gun and operated at 200 kV (Thermo Fisher Scientific) and a Falcon 3 direct electron detector. Tilt series were recorded using Tomography software (Thermo Fisher Scientific) in counting mode and an angular range of – 60° to + 60° with a calibrated physical pixel size of 3.2 Å and a and total dose over the full tilt series of 3.49 e- /Å2 and dose rate 42,16 e-/px/s 3.49 e- /Å2 with 1 second exposure time, 70 um objective aperture. The defocus applied was in a range of −3 µm defocus.

The tomograms were reconstructed using eTomo. Final alignments were done by using 10 nm fiducial gold particles coated with BSA (BSA Gold Tracer, EMS). Gold beads were manually selected and automatically tracked. The fiducial model was corrected in all cases where the automatic tracking failed. Tomograms were binned 2x corresponding to a pixel size of 0.676 nm for the Titan and 0,6368 nm for the Glacios and SIRT-like filter option in eTomo was applied. For visualization purposes, the reconstructed volumes were processed by a Gaussian filter.

### Statistical analysis

All column graphs and statistical analysis were performed by using GraphPad Prism version 7 software. Unpaired t-test was applied to comparisons of two conditions presented in the figure 1 and 4, and in supplementary figures. For more than two groups statistical significance was assessed by a one-way ANOVA with Tukey correction in the figure 5. Quantifications were done blind. Quantitative data depicted as (± SEM) mean standard deviation.

Graph in the Figure 1D showing the percentage of N transfer in co-culture at 24h and 48h. Mean percentage of N transfer in co-culture 24h: 36.47% ± 3.96, co-culture 48h: 62.56% ± 8.28, (p=0.0468 (*) for co-culture 48h versus co-culture 24h; N=3). Graph in the Figure 1G showing the percentage of S transfer in co-culture at 24h and 48h. Mean percentage of S transfer in co-culture 24h: 21.84% ± 5.09, co-culture 48h: 42.44% ± 4.38, (p=0.0374 (*) for co-culture 48h versus co-culture 24h; N=3). Graph in the Figure 6C showing the percentage of TNT connected cells between Vero E6 non-infected and SARS-CoV-2 infected. Mean percentage of TNTs connected Vero non-infected cells: 13.95% ± 2.46. Mean percentage of TNTs connected Vero SARS-CoV-2 infected cells: 44.69% ± 1.96 (p=0.0006 (***) for Vero SARS-CoV-2 versus Vero non-infected; N=3). Graph in the Figure 7C showing the percentage of N transfer in co-culture at 24h and 48h treat and not with the neutralizing antibody. Mean percentage of N transfer in co-culture 24h Control: 95.45% ± 4.29 versus co-culture 24h plus neutralizing antibody 42.91 ± 4.55; p=0.0018 (**) for co-culture 24h Control versus co-culture 24h plus neutralizing antibody. Mean percentage of N transfer in co-culture 48h Control: 96.88% ± 3.12 versus co-culture 48h plus neutralizing antibody 63.90 ± 1.99; p=0.0104 (*) for co-culture 48h Control versus co-culture 48h plus neutralizing antibody. p=0.9914 (ns) for co-culture 24h Control versus co-culture 48h control. p=0.0122 (*) for co-culture 24h Control versus co-culture 48h plus neutralizing antibody. p=0.0016 (**) for co-culture 24h control antibody versus co-culture 48h plus neutralizing antibody. p=0.0496 (*) for co-culture 24h plus neutralizing antibody versus co-culture 48h plus neutralizing antibody. Mean percentage of N transfer in secretion 24h Control: 100% ± 0 versus co-culture 24h plus neutralizing antibody 0 ± 0; p=0.0005 (***) for co-culture 24h Control versus co-culture 24h plus neutralizing antibody. Mean percentage of N transfer in secretion 48h Control: 80% ± 10 versus co-culture 48h plus neutralizing antibody 0 ± 0; p=0.0008 (***) for co-culture 48h Control versus co-culture 48h plus neutralizing antibody. Graph in the Supplementary Figure 4C (left) showing the percentage of N transfer in secretion test at 24h and 48h. Mean percentage of N transfer in secretion 24h: 0% ± 0, co-culture 48h: 1.44% ± 1.44, (p=0.3739 (ns) for secretion 48h versus co-culture 24h; N=3). Graph in the Supplementary Figure 4C (right) showing the percentage of S transfer in secretion test at 24h and 48h. Mean percentage of N transfer in secretion 24h: 0% ± 0, co-culture 48h: 1.44% ± 1.44, (p=0.3739 (ns) for secretion 48h versus co-culture 24h; N=3). Pearson Correlation Coefficient (PCC) was employed to quantify colocalization between anti-S and anti-N. 20 cells were considered. PCC was calculated by using JACoP plugins in Fiji.

## Acknowledgements

The authors thank all the lab members for useful discussion in particular Maura Samarani, Diego Cordero Cervantes and Michael Henderson. The NanoImaging Core at Institut Pasteur is acknowledged for support with image acquisition and analysis. The NanoImaging Core was created with the help of a grant from the French Government’s Investissements d’Avenir program (EQUIPEX CACSICE - Centre d’analyse de systèmes complexes dans les environnements complexes, ANR-11-EQPX-0008). We would like to thank Jean-Marie Winter (NanoImaging Core at Institut Pasteur). We also gratefully acknowledge Anna Sartori-Ruopp (Ultrapole, Institut Pasteur) and Gerard Péhau-Arnaudet.

Nicolas Escriou (Institut Pasteur) for the primary antibody SARS-CoV-anti-N IgG. Cyril Planchais from the group of Hugo Mouquet (Institut Pasteur) for the human SARS-CoV-2 anti-S IgG and neutralizing Antibody (C3-alpha 235). This work was supported by the « URGENCE COVID-19 » fundraising campaign of Institut Pasteur (Paris).

## Author contributions

A.P conceived the experiments and wrote the manuscript; prepared the figures and image rendering; performed co-culture, TNT-counting experiments and all quantifications, prepared and plunch-freeze the cells for TEM experiments, performed all correlative, cryo-correlative light, and electron tomography experiments, tomograms reconstruction. A.P. helped S.P to infect cells. S.P. performed focus forming assay; semi-solid plaque; immunospot and set-up the concentration of the neutralizing antibody. A.P, S.P, M.V, G.B.S, C.Z, discussed the results. All authors commented on the manuscript. C.Z. conceived the project, supervised all the work, and wrote the manuscript. C.Z, M.V, G.B.S. contributed to funding acquisition.

---

**Supplementary Fig. 1.**
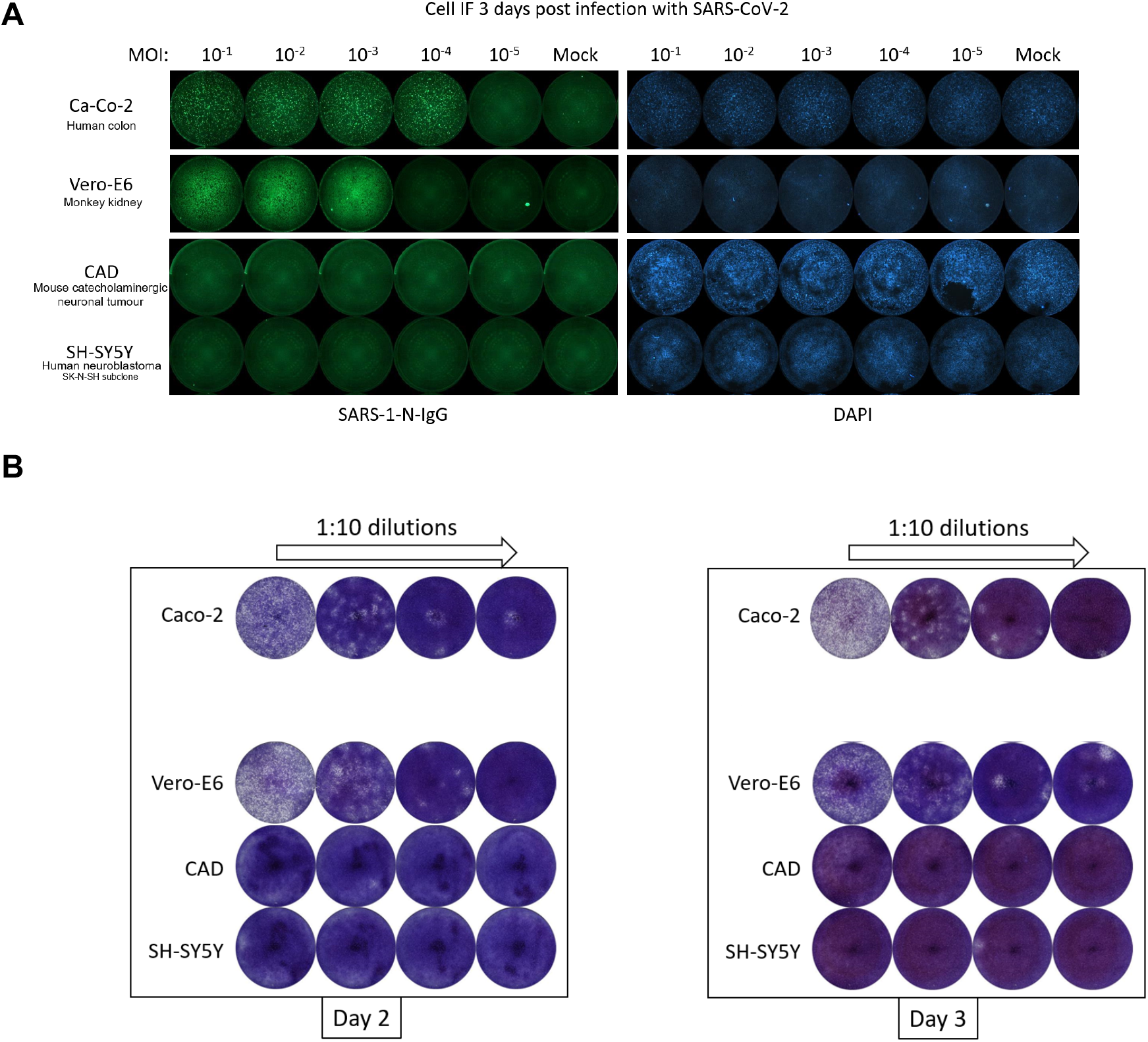
Screening of cell lines for susceptibility to SARS-CoV-2. **(A)** Caco-2, Vero E6, CAD, SH-SY5Y cells were plated on a 96 multi-well plate and infected with SARS-CoV-2 MOI from 10^-1^ to 10^-5^ in DMEM 2% FBS. At 3 days post infection the monolayers were fixed with 4% PFA and viral infection was visualized using an anti-N immunostaining and DAPI to stain the nuclei and then visualized using the Fluoro-X suite of a C.T.L. Immunospot® S6 Image Analyzer. (**B)** At day 2 and 3 post infection of the cells: Caco-2, Vero E6, CAD, SH-SY5Y cells, an aliquot of the supernatant from the higher MOI was collected for titration.

**Supplementary Fig. 2.**
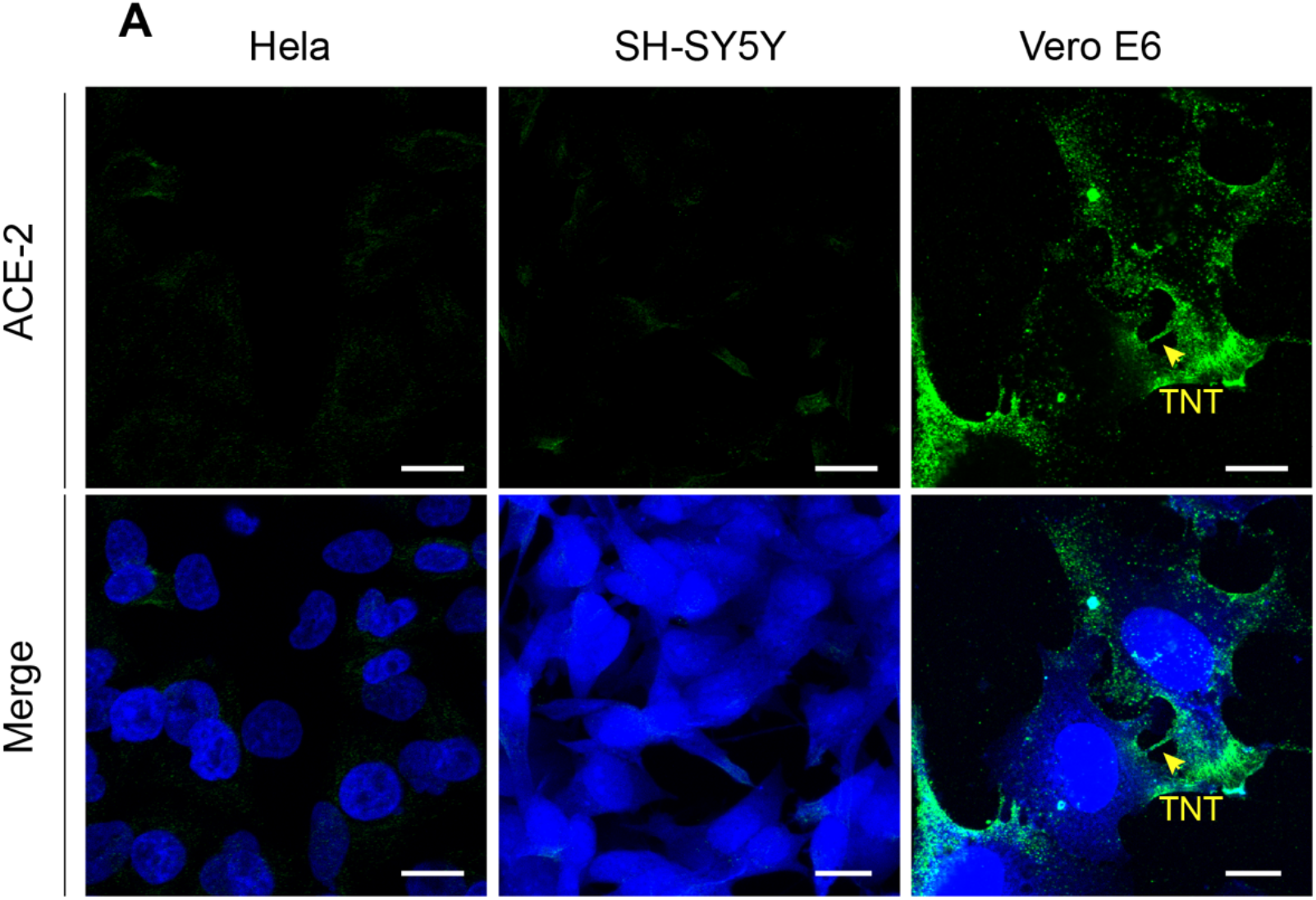
ACE2 receptor expression. **(A)** Confocal micrograph displaying Hela, SH-SY5Y and Vero E6 cells labeled with an anti-ACE2 antibody (green) and cell mask blue. Scale bars: a 15 µm.

**Supplementary Fig. 3.**
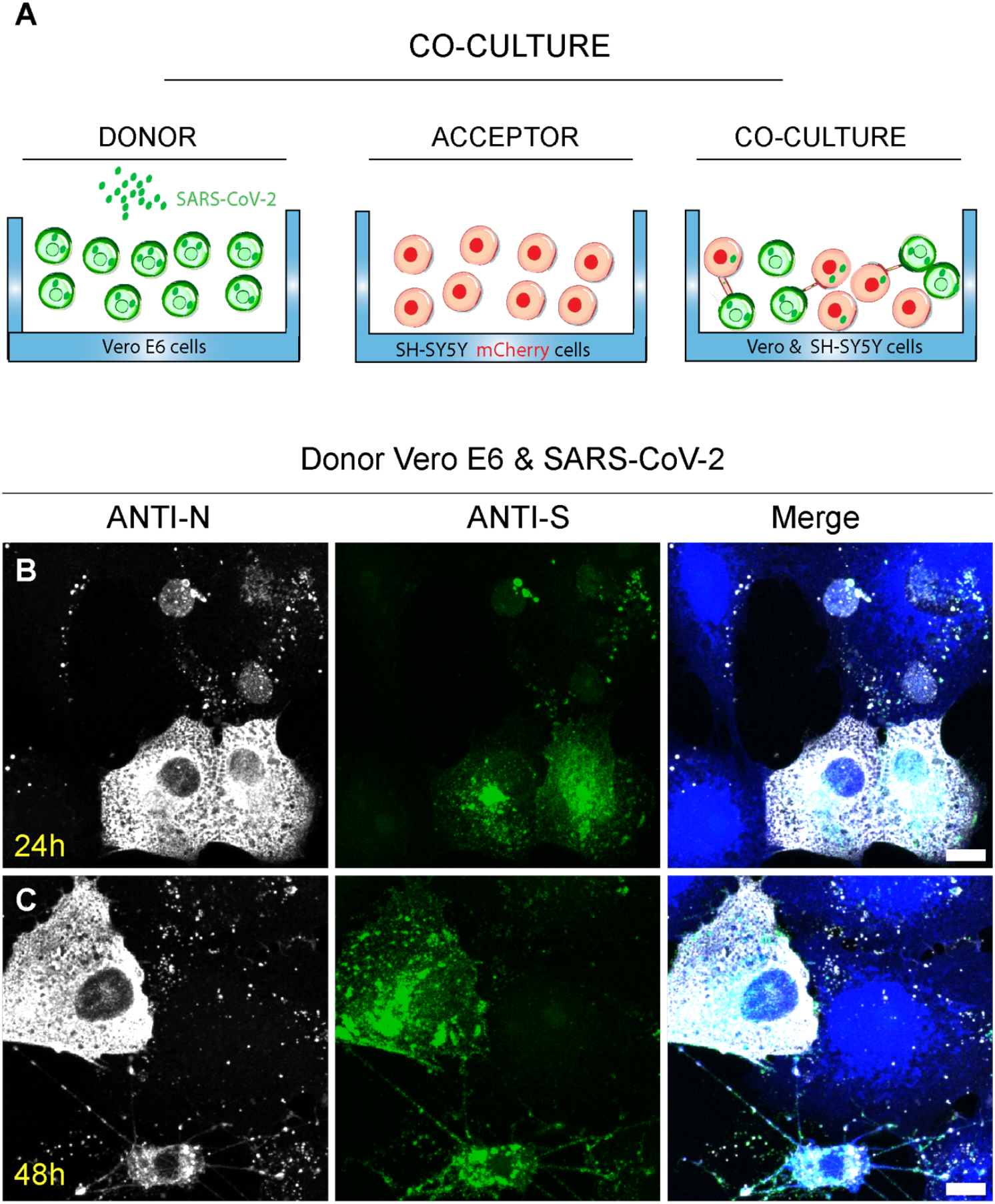
Co-culture pipeline and SARS-CoV-2 detection in Vero E6 donor cells. **(A)** Description of co-culture experiments: Donor Vero E6 cells were infected with SARS-CoV-2 MOI of 0.05 for 48h. After 48h, donor cells were co-cultured with the acceptor SH-SY5Y mCherry cells and incubate for additional 24h and 48h before to be fixed. **(B-C)** Confocal micrographs showing only donor SARS-CoV-2 infected Vero E6 cells used to perform a 24h and 48h co-culture with SH-SY5Y cells (co-culture not showed). Donor cells were fixed at 24h and 48h and immunostained with anti-N and anti-S antibodies to detect SARS-CoV-2. B, C 10µm.

**Supplementary Fig. 4.**
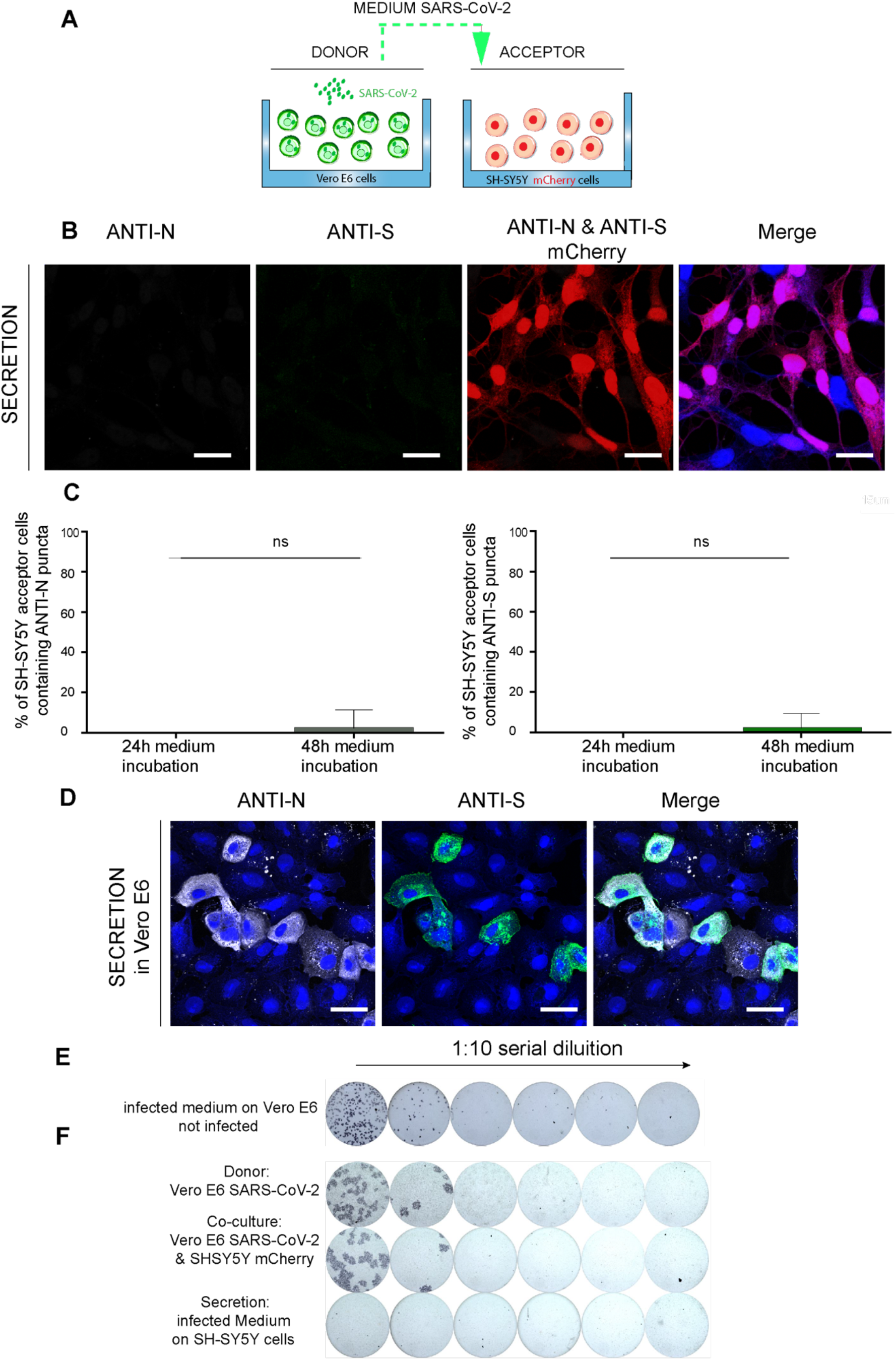
Secretion test. **(A)** Description of secretion experiments: the medium from Vero E6 infected with SARS-CoV-2 MOI of 0.05 was centrifuged and incubated for 24h and 48h with non-permissive SH-SY5Y neuronal cells. (**B)** Confocal micrograph showing SH-SY5Y cells incubated with infected medium from Vero E6 cells. Cells were fixed after 48h of incubation at 37° and 5% CO2 and immunostained with an anti-N antibody and anti-S antibody to detect SARS-CoV-2. (**C)** Left graph showing the percentage of N transfer in secretion test at 24h and 48h. Mean percentage of N transfer in secretion 24h: 0% ± 0, co-culture 48h: 1.44% ± 1.44, (p=0.3739 (ns) for secretion 48h versus co-culture 24h; N=3). Right graph showing the percentage of S transfer in secretion test at 24h and 48h. Mean percentage of S transfer in secretion 24h: 0% ± 0, co-culture 48h: 1.44% ± 1.44, (p=0.3739 (ns) for secretion 48h versus co-culture 24h; N=3). (**D)** Confocal micrograph showing not-infected Vero E6 cells incubated with infected medium from SARS-CoV-2 infected Vero E6 cells. Cells were fixed after 48h of incubation at 37° and 5% CO2 and immunostained with an anti-N antibody and anti-S antibody to detect SARS-CoV-2 particles. (**E)** The infectious titer of the supernatant used to infect Vero E6 cells was calculated using a focus forming assay. (**F)** The 48h supernatants from donor infected cells, from co-culture and from secretion test were used to assess viral production by focus forming assay titration protocol. Scale bars: **B, D** 20 µm.

**Supplementary Fig. 5.**
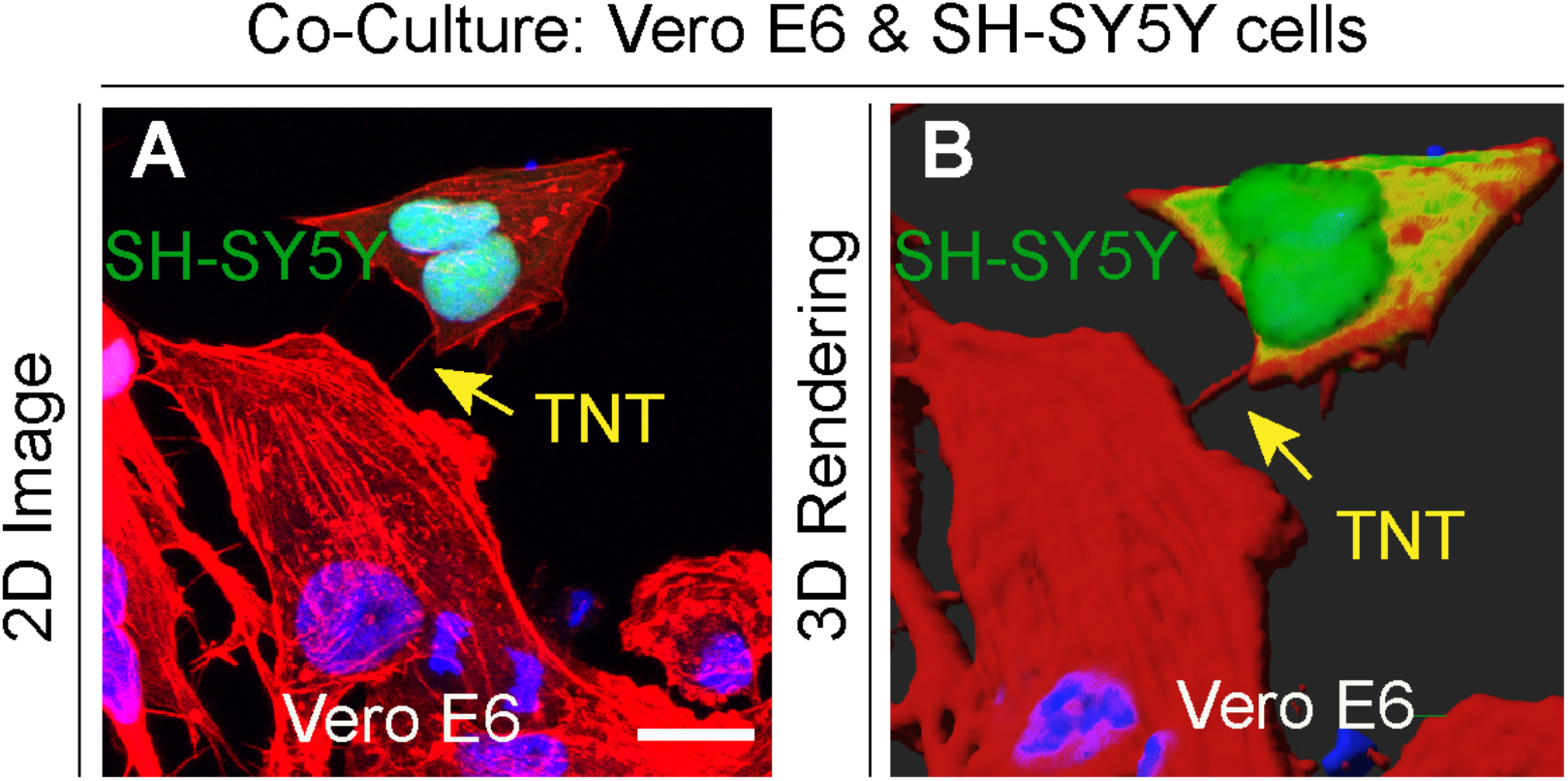
TNTs are formed between Vero E6 cells and SH-SY5Y cells. **(A)** Representative confocal micrograph displaying TNT between Vero E6 cells and GFP NLS SH-SY5Y cells. Cells were stained with Phalloidin Rhodamine (red) to label the plasma membrane and TNTs, and DAPI (blue) to label the nuclei. (**B)** 3D rendering of (**A**). Scale bars: **A**, 15 µm.

**Supplementary Fig. 6.**
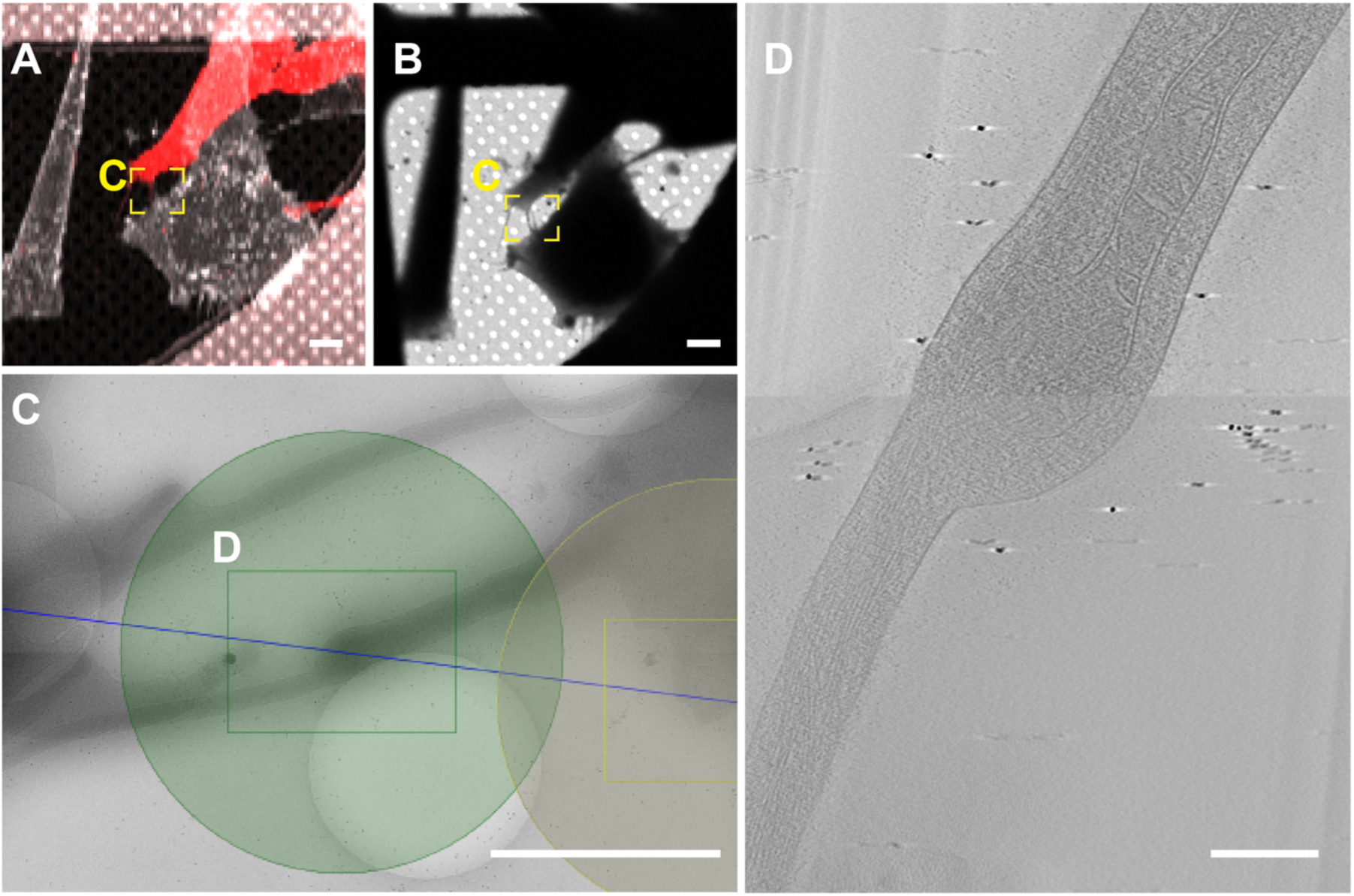
Correlative Cryo-EM on TNT-connected not-infected Vero E6 and SH-SY5Y mCherry cells. (**A)** Confocal micrograph shows TNT-connected naïve Vero E6 cells and SH-SY5Y mCherry cells. **(B** and **C)** Low magnification (B) and intermediate magnification (C) of electron micrograph of TNT-connected naïve Vero E6 cells and SH-SY5Y mCherry cells. **(D)** High magnification cryo-ET slice corresponding to the green rectangle in (C). Scale bars, A, B 10µm; C, 2 µm; D, 100 nm.

**Supplementary Fig. 7.**
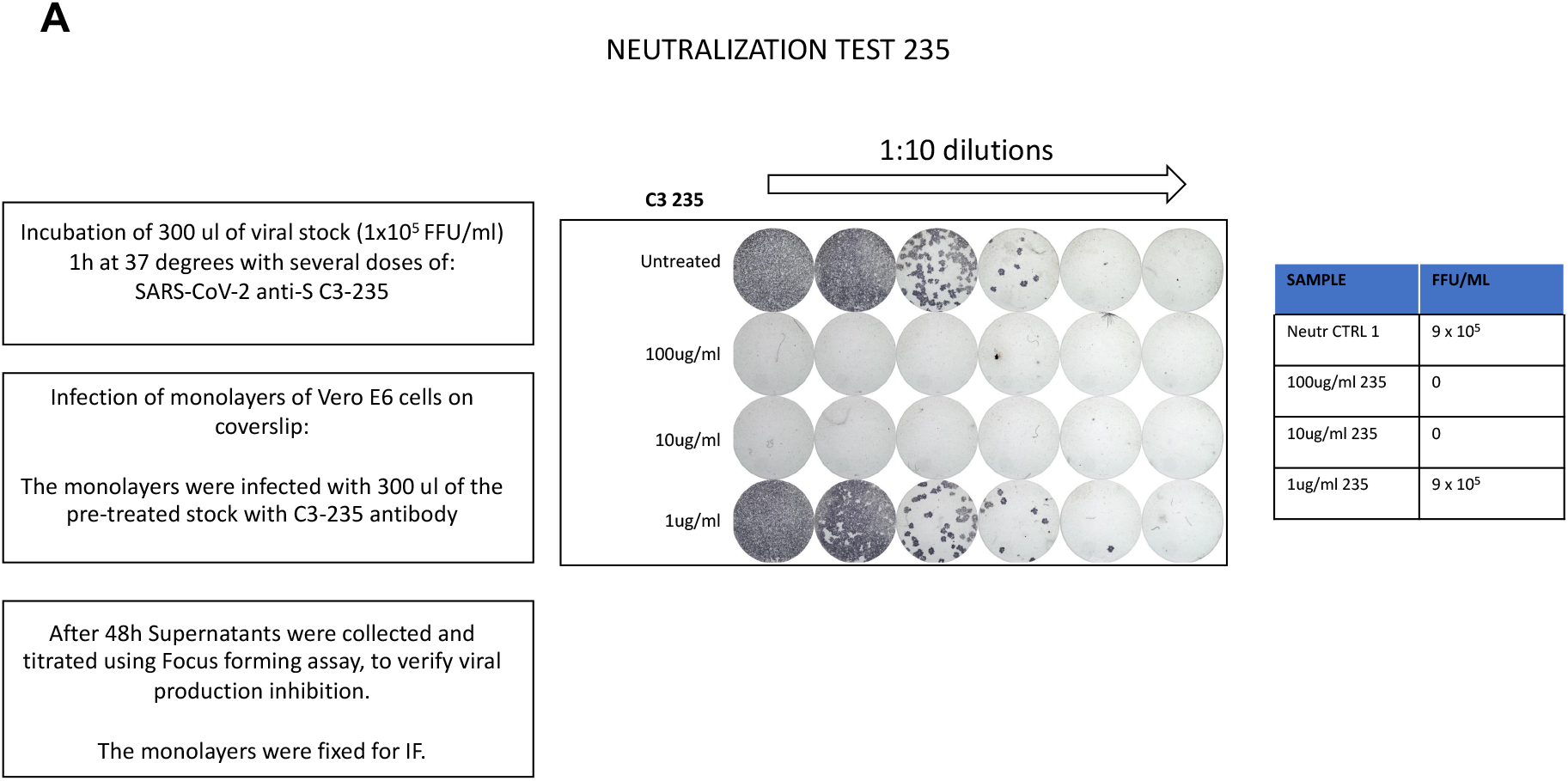
Experimental set-up to identify the minimal concentration of SARS-CoV-2 IgG C3 235 sufficient to have a complete neutralization of the viral stock of 1-5 x 10^5^ FFU/ml. **(A)** Three different concentrations of human SARS-CoV-2 IgG C3 235 (1, 10 and 100 ug/ml) were incubated 1h at 37°C, 5% CO2 and then used to infect monolayers of Vero E6 cells for 48h. Viral production was then assessed directly by processing the monolayers and titration of the supernatant by focus forming assay.

**Supplementary Fig. 8.**
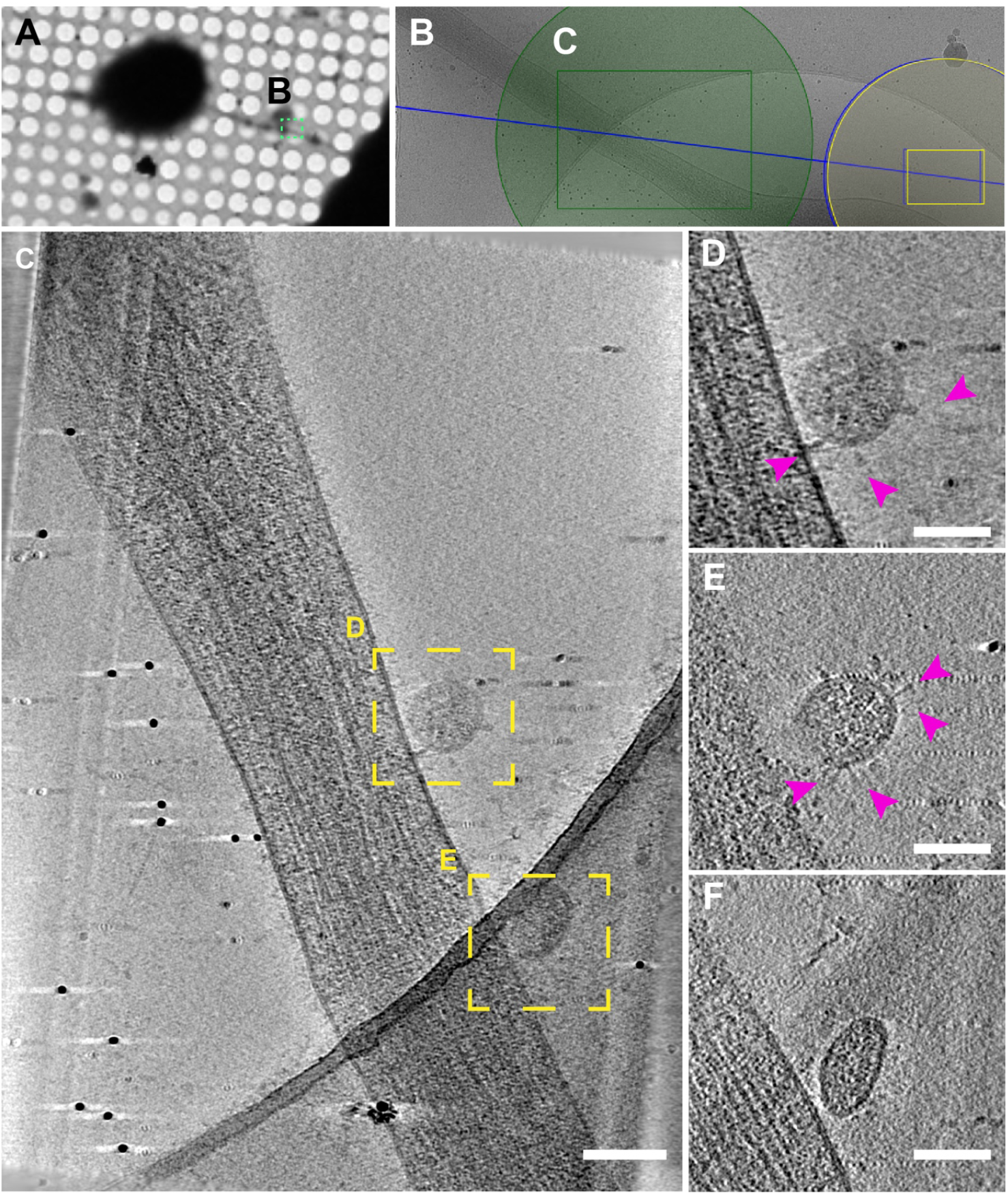
SARS-CoV-2 on TNTs connected Vero E6 infected cells. (**A** and **B)** Low (A) and intermediate (B) magnification of an electron micrograph of TNT-connected SARS-CoV-2 infected Vero E6 cells. (**C)** High magnification cryo-ET slice corresponding to the green rectangle in (B). (**D**, **E** and **F**) High-magnification cryo-tomography slices showing the extracellular virions on TNT. Pink arrowheads indicate the spike on SARS-CoV2. Scale bars, (A), 10µm; (B), 1 µm; (C), 150 nm, (D), (E), (F) 100 nm.

## Description of Supplementary Files

**File Name:** Supplementary Movie 1

**Description: Description:** Representative slices of a reconstructed tomogram displaying TNT between SARS-CoV-2 Vero E6 infected cells and mCherry SH-SY5Y cells shown in Fig. 5D. These slices display Double Membrane Vesicles (DMV) inside TNTs. Scale bar: 200nm.

**File Name:** Supplementary Movie 2

**Description: Description:** Representative slices of a reconstructed tomogram displaying TNT between SARS-CoV-2 Vero E6 infected cells and mCherry SH-SY5Y cells shown in Fig. 5I. These slices display several vesicular compartments inside TNTs. Scale bar: 100nm.

**File Name:** Supplementary Movie 3

**Description:** Representative slices of a reconstructed tomogram displaying TNT between not-infected Vero E6 cells and mCherry SH-SY5Y cells shown in Supplementary Fig. 6D. These slices display mitochondria inside TNTs. Scale bar: 100nm.

**File Name:** Supplementary Movie 4

**Description:** Representative slices of a reconstructed tomogram displaying TNT connecting two SARS-CoV-2-infected Vero E6 cells shown in Fig. 8 Scale bar: 200nm.

**File Name:** Supplementary Movie 5

**Description:** Representative slices of a reconstructed tomogram displaying TNT arise from SARS-CoV-2 Vero E6 infected cells positive for the anti-Spike antibody containing vesicles compartments and SARS-CoV-2 like structure on the surface. Scale bar: 200nm

**File Name:** Supplementary Movie 6

**Description:** Representative slices of a reconstructed tomogram displaying TNT arise from SARS-CoV-2-infected Vero E6 cells show in Supplementary Fig. 8. Scale bar: 200nm

## References

1. Zhou P, Yang X-L, Wang X-G, Hu B, Zhang L, Zhang W, et al. A pneumonia outbreak associated with a new coronavirus of probable bat origin. Nature [Internet]. 2020 Mar [cited 2021 Jul 21];579(7798):270–3. Available from: https://www.nature.com/articles/s41586-020-2012-7

2. Huang N, Pérez P, Kato T, Mikami Y, Okuda K, Gilmore RC, et al. SARS-CoV-2 infection of the oral cavity and saliva. Nat Med [Internet]. 2021 May [cited 2021 Jul 21];27(5):892–903. Available from: https://www.nature.com/articles/s41591-021-01296-8

3. Zhu N, Zhang D, Wang W, Li X, Yang B, Song J, et al. A Novel Coronavirus from Patients with Pneumonia in China, 2019. N Engl J Med [Internet]. 2020 Feb 20 [cited 2021 Jul 21];382(8):727–33. Available from: https://doi.org/10.1056/NEJMoa2001017

4. Mao L, Jin H, Wang M, Hu Y, Chen S, He Q, et al. Neurologic Manifestations of Hospitalized Patients With Coronavirus Disease 2019 in Wuhan, China. JAMA Neurol. 2020 Jun 1;77(6):683–90.

5. Song E, Zhang C, Israelow B, Lu-Culligan A, Prado AV, Skriabine S, et al. Neuroinvasion of SARS-CoV-2 in human and mouse brain. J Exp Med. 2021 Mar 1;218(3):e20202135.

6. Huang C, Wang Y, Li X, Ren L, Zhao J, Hu Y, et al. Clinical features of patients infected with 2019 novel coronavirus in Wuhan, China. The Lancet [Internet]. 2020 Feb 15 [cited 2021 Jul 21];395(10223):497–506. Available from: https://www.thelancet.com/journals/lancet/article/PIIS0140-6736(20)30183-5/abstract

7. Conde Cardona G, Quintana Pájaro LD, Quintero Marzola ID, Ramos Villegas Y, Moscote Salazar LR. Neurotropism of SARS-CoV 2: Mechanisms and manifestations. J Neurol Sci. 2020 May 15;412:116824.

8. Davis HE, Assaf GS, McCorkell L, Wei H, Low RJ, Re’em Y, et al. Characterizing Long COVID in an International Cohort: 7 Months of Symptoms and Their Impact. medRxiv [Internet]. 2020 Dec 27 [cited 2021 Jul 21];2020.12.24.20248802. Available from: https://www.medrxiv.org/content/10.1101/2020.12.24.20248802v2

9. Taribagil P, Creer D, Tahir H. ‘Long COVID’ syndrome. BMJ Case Rep CP [Internet]. 2021 Apr 1 [cited 2021 Jul 21];14(4):e241485. Available from: https://casereports.bmj.com/content/14/4/e241485

10. Ziauddeen N, Gurdasani D, O’Hara ME, Hastie C, Roderick P, Yao G, et al. Characteristics of Long Covid: findings from a social media survey. medRxiv [Internet]. 2021 Mar 27 [cited 2021 Jul 21];2021.03.21.21253968. Available from: https://www.medrxiv.org/content/10.1101/2021.03.21.21253968v2

11. Zubair AS, McAlpine LS, Gardin T, Farhadian S, Kuruvilla DE, Spudich S. Neuropathogenesis and Neurologic Manifestations of the Coronaviruses in the Age of Coronavirus Disease 2019: A Review. JAMA Neurol. 2020 Aug 1;77(8):1018–27.

12. Cyranoski D. Profile of a killer: the complex biology powering the coronavirus pandemic. Nature [Internet]. 2020 May 4 [cited 2021 Jul 21];581(7806):22–6. Available from: https://www.nature.com/articles/d41586-020-01315-7

13. Matschke J, Lütgehetmann M, Hagel C, Sperhake JP, Schröder AS, Edler C, et al. Neuropathology of patients with COVID-19 in Germany: a post-mortem case series. Lancet Neurol [Internet]. 2020 Nov 1 [cited 2021 Jul 21];19(11):919–29. Available from: https://www.thelancet.com/journals/laneur/article/PIIS1474-4422(20)30308-2/abstract

14. Bullen CK, Hogberg HT, Bahadirli-Talbott A, Bishai WR, Hartung T, Keuthan C, et al. Infectability of human BrainSphere neurons suggests neurotropism of SARS-CoV-2. ALTEX. 2020;37(4):665–71.

15. Jacob F, Pather SR, Huang W-K, Wong SZH, Zhou H, Zhang F, et al. Human Pluripotent Stem Cell-Derived Neural Cells and Brain Organoids Reveal SARS-CoV-2 Neurotropism. bioRxiv [Internet]. 2020 Jul 28 [cited 2021 Jul 21];2020.07.28.225151. Available from: https://www.biorxiv.org/content/10.1101/2020.07.28.225151v1

16. Pellegrini L, Albecka A, Mallery DL, Kellner MJ, Paul D, Carter AP, et al. SARS-CoV-2 Infects the Brain Choroid Plexus and Disrupts the Blood-CSF Barrier in Human Brain Organoids. Cell Stem Cell [Internet]. 2020 Dec 3 [cited 2021 Jul 21];27(6):951–961.e5. Available from: https://www.ncbi.nlm.nih.gov/pmc/articles/PMC7553118/

17. Puelles VG, Lütgehetmann M, Lindenmeyer MT, Sperhake JP, Wong MN, Allweiss L, et al. Multiorgan and Renal Tropism of SARS-CoV-2. N Engl J Med. 2020 Aug 6;383(6):590–2.

18. Moriguchi T, Harii N, Goto J, Harada D, Sugawara H, Takamino J, et al. A first case of meningitis/encephalitis associated with SARS-Coronavirus-2. Int J Infect Dis IJID Off Publ Int Soc Infect Dis. 2020 May;94:55–8.

19. Meinhardt J, Radke J, Dittmayer C, Franz J, Thomas C, Mothes R, et al. Olfactory transmucosal SARS-CoV-2 invasion as a port of central nervous system entry in individuals with COVID-19. Nat Neurosci [Internet]. 2021 Feb [cited 2021 Jul 21];24(2):168–75. Available from: https://www.nature.com/articles/s41593-020-00758-5

20. Sun S-H, Chen Q, Gu H-J, Yang G, Wang Y-X, Huang X-Y, et al. A Mouse Model of SARS-CoV-2 Infection and Pathogenesis. Cell Host Microbe. 2020 Jul 8;28(1):124–133.e4.

21. Jha NK, Ojha S, Jha SK, Dureja H, Singh SK, Shukla SD, et al. Evidence of Coronavirus (CoV) Pathogenesis and Emerging Pathogen SARS-CoV-2 in the Nervous System: A Review on Neurological Impairments and Manifestations. J Mol Neurosci [Internet]. 2021 Jan 19 [cited 2021 Jul 21]; Available from: https://doi.org/10.1007/s12031-020-01767-6

22. Pacheco-Herrero M, Soto-Rojas LO, Harrington CR, Flores-Martinez YM, Villegas-Rojas MM, León-Aguilar AM, et al. Elucidating the Neuropathologic Mechanisms of SARS-CoV-2 Infection. Front Neurol [Internet]. 2021 [cited 2021 Jul 21];0. Available from: https://www.frontiersin.org/articles/10.3389/fneur.2021.660087/full

23. Andersen KG, Rambaut A, Lipkin WI, Holmes EC, Garry RF. The proximal origin of SARS-CoV-2. Nat Med [Internet]. 2020 Apr [cited 2021 Jul 21];26(4):450–2. Available from: https://www.nature.com/articles/s41591-020-0820-9

24. Hoffmann M, Kleine-Weber H, Schroeder S, Krüger N, Herrler T, Erichsen S, et al. SARS-CoV-2 Cell Entry Depends on ACE2 and TMPRSS2 and Is Blocked by a Clinically Proven Protease Inhibitor. Cell [Internet]. 2020 Apr 16 [cited 2021 Jul 21];181(2):271–280.e8. Available from: https://www.sciencedirect.com/science/article/pii/S0092867420302294

25. Jackson CB, Farzan M, Chen B, Choe H. Mechanisms of SARS-CoV-2 entry into cells. Nat Rev Mol Cell Biol [Internet]. 2021 Oct 5 [cited 2021 Nov 2];1–18. Available from: https://www.nature.com/articles/s41580-021-00418-x

26. Chen R, Wang K, Yu J, Howard D, French L, Chen Z, et al. The Spatial and Cell-Type Distribution of SARS-CoV-2 Receptor ACE2 in the Human and Mouse Brains. Front Neurol [Internet]. 2021 Jan 20 [cited 2021 Oct 5];11:573095. Available from: https://www.ncbi.nlm.nih.gov/pmc/articles/PMC7855591/

27. Serrano GE, Walker JE, Arce R, Glass MJ, Vargas D, Sue LI, et al. Mapping of SARS-CoV-2 Brain Invasion and Histopathology in COVID-19 Disease. medRxiv [Internet]. 2021 Feb 18 [cited 2021 Jul 21];2021.02.15.21251511. Available from: https://www.medrxiv.org/content/10.1101/2021.02.15.21251511v1

28. Victoria GS, Zurzolo C. The spread of prion-like proteins by lysosomes and tunneling nanotubes: Implications for neurodegenerative diseases. J Cell Biol [Internet]. 2017 Jul 19 [cited 2021 Jul 21];216(9):2633–44. Available from: https://doi.org/10.1083/jcb.201701047

29. Hawkes CH, Del Tredici K, Braak H. Parkinson’s disease: the dual hit theory revisited. Ann N Y Acad Sci. 2009 Jul;1170:615–22.

30. Gousset K, Schiff E, Langevin C, Marijanovic Z, Caputo A, Browman DT, et al. Prions hijack tunnelling nanotubes for intercellular spread. Nat Cell Biol. 2009 Mar;11(3):328–36.

31. Abounit S, Bousset L, Loria F, Zhu S, de Chaumont F, Pieri L, et al. Tunneling nanotubes spread fibrillar α-synuclein by intercellular trafficking of lysosomes. EMBO J. 2016 Oct 4;35(19):2120–38.

32. Vargas JY, Loria F, Wu Y-J, Córdova G, Nonaka T, Bellow S, et al. The Wnt/Ca2+ pathway is involved in interneuronal communication mediated by tunneling nanotubes. EMBO J. 2019 Dec 2;38(23):e101230.

33. Rustom A, Saffrich R, Markovic I, Walther P, Gerdes H-H. Nanotubular highways for intercellular organelle transport. Science. 2004 Feb 13;303(5660):1007–10.

34. Cordero Cervantes D, Zurzolo C. Peering into tunneling nanotubes-The path forward. EMBO J. 2021 Apr 15;40(8):e105789.

35. Sartori-Rupp A, Cordero Cervantes D, Pepe A, Gousset K, Delage E, Corroyer-Dulmont S, et al. Correlative cryo-electron microscopy reveals the structure of TNTs in neuronal cells. Nat Commun [Internet]. 2019 Jan 21 [cited 2021 Jul 21];10(1):342. Available from: https://www.nature.com/articles/s41467-018-08178-7

36. Abounit S, Zurzolo C. Wiring through tunneling nanotubes--from electrical signals to organelle transfer. J Cell Sci. 2012 Mar 1;125(Pt 5):1089–98.

37. Gerdes H-H, Carvalho RN. Intercellular transfer mediated by tunneling nanotubes. Curr Opin Cell Biol. 2008 Aug;20(4):470–5.

38. Eugenin EA, Gaskill PJ, Berman JW. Tunneling nanotubes (TNT) are induced by HIV-infection of macrophages: a potential mechanism for intercellular HIV trafficking. Cell Immunol. 2009;254(2):142–8.

39. Jansens RJJ, Tishchenko A, Favoreel HW. Bridging the Gap: Virus Long-Distance Spread via Tunneling Nanotubes. J Virol. 2020 Mar 31;94(8):e02120–19.

40. Kadiu I, Gendelman HE. Human Immunodeficiency Virus type 1 Endocytic Trafficking Through Macrophage Bridging Conduits Facilitates Spread of Infection. J Neuroimmune Pharmacol [Internet]. 2011 [cited 2021 Jul 21];6(4):658–75. Available from: https://www.ncbi.nlm.nih.gov/pmc/articles/PMC3232570/

41. Souriant S, Balboa L, Dupont M, Pingris K, Kviatcovsky D, Cougoule C, et al. Tuberculosis Exacerbates HIV-1 Infection through IL-10/STAT3-Dependent Tunneling Nanotube Formation in Macrophages. Cell Rep. 2019 Mar 26;26(13):3586–3599.e7.

42. Sowinski S, Jolly C, Berninghausen O, Purbhoo MA, Chauveau A, Köhler K, et al. Membrane nanotubes physically connect T cells over long distances presenting a novel route for HIV-1 transmission. Nat Cell Biol. 2008 Feb;10(2):211–9.

43. Wan Y, Shang J, Graham R, Baric RS, Li F. Receptor Recognition by the Novel Coronavirus from Wuhan: an Analysis Based on Decade-Long Structural Studies of SARS Coronavirus. J Virol. 2020 Mar 17;94(7):e00127–20.

44. Ge X-Y, Li J-L, Yang X-L, Chmura AA, Zhu G, Epstein JH, et al. Isolation and characterization of a bat SARS-like coronavirus that uses the ACE2 receptor. Nature [Internet]. 2013 Nov [cited 2021 Jul 22];503(7477):535–8. Available from: https://www.nature.com/articles/nature12711

45. Li Y, Bai W, Hashikawa T. The neuroinvasive potential of SARS-CoV2 may play a role in the respiratory failure of COVID-19 patients. J Med Virol [Internet]. 2020 Mar 11 [cited 2021 Jul 22];10.1002/jmv.25728. Available from: https://www.ncbi.nlm.nih.gov/pmc/articles/PMC7228394/

46. Ogando NS, Dalebout TJ, Zevenhoven-Dobbe JC, Limpens RWAL, van der Meer Y, Caly L, et al. SARS-coronavirus-2 replication in Vero E6 cells: replication kinetics, rapid adaptation and cytopathology. J Gen Virol. 2020 Sep;101(9):925–40.

47. Matheson NJ, Lehner PJ. How does SARS-CoV-2 cause COVID-19? Science. 2020 Jul 31;369(6503):510–1.

48. Chen N, Zhou M, Dong X, Qu J, Gong F, Han Y, et al. Epidemiological and clinical characteristics of 99 cases of 2019 novel coronavirus pneumonia in Wuhan, China: a descriptive study. The Lancet [Internet]. 2020 Feb 15 [cited 2021 Jul 22];395(10223):507–13. Available from: https://www.thelancet.com/journals/lancet/article/PIIS0140-6736(20)30211-7/abstract

49. Helms J, Kremer S, Merdji H, Clere-Jehl R, Schenck M, Kummerlen C, et al. Neurologic Features in Severe SARS-CoV-2 Infection. N Engl J Med. 2020 Jun 4;382(23):2268–70.

50. Poyiadji N, Shahin G, Noujaim D, Stone M, Patel S, Griffith B. COVID-19-associated Acute Hemorrhagic Necrotizing Encephalopathy: Imaging Features. Radiology. 2020 Aug;296(2):E119–20.

51. Sedaghat Z, Karimi N. Guillain Barre syndrome associated with COVID-19 infection: A case report. J Clin Neurosci Off J Neurosurg Soc Australas. 2020 Jun;76:233–5.

52. Virani A, Rabold E, Hanson T, Haag A, Elrufay R, Cheema T, et al. Guillain-Barré Syndrome associated with SARS-CoV-2 infection. IDCases. 2020;20:e00771.

53. Coolen T, Lolli V, Sadeghi N, Rovai A, Trotta N, Taccone FS, et al. Early postmortem brain MRI findings in COVID-19 non-survivors. Neurology. 2020 Oct 6;95(14):e2016–27.

54. Baig AM. Neurological manifestations in COVID-19 caused by SARS-CoV-2. CNS Neurosci Ther [Internet]. 2020 [cited 2021 Jul 22];26(5):499–501. Available from: https://onlinelibrary.wiley.com/doi/abs/10.1111/cns.13372

55. De Felice FG, Tovar-Moll F, Moll J, Munoz DP, Ferreira ST. Severe Acute Respiratory Syndrome Coronavirus 2 (SARS-CoV-2) and the Central Nervous System. Trends Neurosci. 2020 Jun;43(6):355–7.

56. V’kovski P, Kratzel A, Steiner S, Stalder H, Thiel V. Coronavirus biology and replication: implications for SARS-CoV-2. Nat Rev Microbiol [Internet]. 2021 Mar [cited 2021 Oct 5];19(3):155–70. Available from: https://www.nature.com/articles/s41579-020-00468-6

57. Son K-N, Liang Z, Lipton HL. Double-Stranded RNA Is Detected by Immunofluorescence Analysis in RNA and DNA Virus Infections, Including Those by Negative-Stranded RNA Viruses. J Virol [Internet]. 2015 Aug 19 [cited 2021 Jul 22];89(18):9383–92. Available from: https://www.ncbi.nlm.nih.gov/pmc/articles/PMC4542381/

58. Klein S, Cortese M, Winter SL, Wachsmuth-Melm M, Neufeldt CJ, Cerikan B, et al. SARS-CoV-2 structure and replication characterized by in situ cryo-electron tomography. Nat Commun [Internet]. 2020 Nov 18 [cited 2021 Jul 22];11(1):5885. Available from: https://www.nature.com/articles/s41467-020-19619-7

59. Wolff G, Limpens RWAL, Zevenhoven-Dobbe JC, Laugks U, Zheng S, Jong AWM de, et al. A molecular pore spans the double membrane of the coronavirus replication organelle. Science [Internet]. 2020 Sep 11 [cited 2021 Jul 22];369(6509):1395–8. Available from: https://science.sciencemag.org/content/369/6509/1395

60. Knoops K, Kikkert M, Worm SHE van den, Zevenhoven-Dobbe JC, Meer Y van der, Koster AJ, et al. SARS-Coronavirus Replication Is Supported by a Reticulovesicular Network of Modified Endoplasmic Reticulum. PLOS Biol [Internet]. 2008 Sep 16 [cited 2021 Jul 22];6(9):e226. Available from: https://journals.plos.org/plosbiology/article?id=10.1371/journal.pbio.0060226

61. Snijder EJ, Limpens RWAL, de Wilde AH, de Jong AWM, Zevenhoven-Dobbe JC, Maier HJ, et al. A unifying structural and functional model of the coronavirus replication organelle: Tracking down RNA synthesis. PLoS Biol. 2020 Jun;18(6):e3000715.

62. Maier HJ, Hawes PC, Cottam EM, Mantell J, Verkade P, Monaghan P, et al. Infectious bronchitis virus generates spherules from zippered endoplasmic reticulum membranes. mBio. 2013 Oct 22;4(5):e00801–00813.

63. Ulasli M, Verheije MH, de Haan CAM, Reggiori F. Qualitative and quantitative ultrastructural analysis of the membrane rearrangements induced by coronavirus. Cell Microbiol. 2010 Jun;12(6):844–61.

64. de Haan CAM, Rottier PJM. Molecular interactions in the assembly of coronaviruses. Adv Virus Res. 2005;64:165–230.

65. Delage E, Cervantes DC, Pénard E, Schmitt C, Syan S, Disanza A, et al. Differential identity of Filopodia and Tunneling Nanotubes revealed by the opposite functions of actin regulatory complexes. Sci Rep [Internet]. 2016 Dec 23 [cited 2021 Jul 22];6(1):39632. Available from: https://www.nature.com/articles/srep39632

66. Abounit S, Delage E, Zurzolo C. Identification and Characterization of Tunneling Nanotubes for Intercellular Trafficking. Curr Protoc Cell Biol. 2015 Jun 1;67:12.10.1–12.10.21.

67. Song W, Gui M, Wang X, Xiang Y. Cryo-EM structure of the SARS coronavirus spike glycoprotein in complex with its host cell receptor ACE2. PLOS Pathog [Internet]. 2018 Aug 13 [cited 2021 Jul 22];14(8):e1007236. Available from: https://journals.plos.org/plospathogens/article?id=10.1371/journal.ppat.1007236

68. Liu C, Mendonça L, Yang Y, Gao Y, Shen C, Liu J, et al. The Architecture of Inactivated SARS-CoV-2 with Postfusion Spikes Revealed by Cryo-EM and Cryo-ET. Structure [Internet]. 2020 Nov 3 [cited 2021 Jul 22];28(11):1218–1224.e4. Available from: https://www.sciencedirect.com/science/article/pii/S0969212620303725

69. Yao H, Song Y, Chen Y, Wu N, Xu J, Sun C, et al. Molecular Architecture of the SARS-CoV-2 Virus. Cell [Internet]. 2020 Oct 29 [cited 2021 Jul 22];183(3):730–738.e13. Available from: https://www.sciencedirect.com/science/article/pii/S0092867420311594

70. Caldas LA, Carneiro FA, Higa LM, Monteiro FL, da Silva GP, da Costa LJ, et al. Ultrastructural analysis of SARS-CoV-2 interactions with the host cell via high resolution scanning electron microscopy. Sci Rep [Internet]. 2020 Dec [cited 2021 Oct 5];10(1):16099. Available from: https://www.nature.com/articles/s41598-020-73162-5

71. Lehmann MJ, Sherer NM, Marks CB, Pypaert M, Mothes W. Actin-and myosin-driven movement of viruses along filopodia precedes their entry into cells. J Cell Biol. 2005 Jul 18;170(2):317–25.

72. Najjar FE, Cifuentes-Muñoz N, Chen J, Zhu H, Buchholz UJ, Moncman CL, et al. Human metapneumovirus Induces Reorganization of the Actin Cytoskeleton for Direct Cell-to-Cell Spread. PLOS Pathog [Internet]. 2016 Sep 28 [cited 2021 Oct 5];12(9):e1005922. Available from: https://journals.plos.org/plospathogens/article?id=10.1371/journal.ppat.1005922

73. Bouhaddou M, Memon D, Meyer B, White KM, Rezelj VV, Correa Marrero M, et al. The Global Phosphorylation Landscape of SARS-CoV-2 Infection. Cell [Internet]. 2020 Aug [cited 2021 Oct 5];182(3):685–712.e19. Available from: https://linkinghub.elsevier.com/retrieve/pii/S0092867420308114

74. Dilsizoglu Senol A, Pepe A, Grudina C, Sassoon N, Reiko U, Bousset L, et al. Effect of tolytoxin on tunneling nanotube formation and function. Sci Rep. 2019 Apr 5;9(1):5741.

75. Hagen WJH, Wan W, Briggs JAG. Implementation of a cryo-electron tomography tilt-scheme optimized for high resolution subtomogram averaging. J Struct Biol [Internet]. 2017 Feb 1 [cited 2021 Jul 22];197(2):191–8. Available from: https://www.sciencedirect.com/science/article/pii/S1047847716301137

